# G6PD Maintains Redox Homeostasis and Biosynthesis in LKB1-Deficient KRAS-Driven Lung Cancer

**DOI:** 10.1101/2023.10.06.561131

**Authors:** Taijin Lan, Sara Arastu, Samuel Wang, Jarrick Lam, Wenping Wang, Vrushank Bhatt, Eduardo Cararo Lopes, Zhixian Hu, Michael Sun, Xuefei Luo, Jonathan M. Ghergurovich, Changlong Li, Xiaoyang Su, Joshua D. Rabinowitz, Eileen White, Jessie Yanxiang Guo

**Affiliations:** Rutgers Cancer Institute of New Jersey, New Brunswick, New Jersey 08901, USA; West China School of Basic Medical Sciences & Forensic Medicine, Sichuan University, Chengdu 610041, China; Department of Molecular Biology, Princeton University, Princeton, New Jersey 08544, USA; Department of Chemistry, Princeton University, Princeton, New Jersey 08544, USA; Department of Medicine, Rutgers Robert Wood Johnson Medical School, New Brunswick, New Jersey 08901, USA; Ludwig Princeton Branch, Ludwig Institute for Cancer Research, Princeton University, Princeton, New Jersey 08540, USA; Department of Molecular Biology and Biochemistry, Rutgers University, Piscataway, New Jersey 08854, USA; Department of Chemical Biology, Rutgers Ernest Mario School of Pharmacy, Piscataway, New Jersey 08854, USA

**Keywords:** G6PD, KRAS, LKB1, NSCLC, NADPH, Redox, fatty acid synthesis, metabolic reprogramming, serine metabolism

## Abstract

Cancer cells depend on nicotinamide adenine dinucleotide phosphate (NADPH) to combat oxidative stress and support reductive biosynthesis. One major NAPDH production route is the oxidative pentose phosphate pathway (committed step: glucose-6-phosphate dehydrogenase, G6PD). Alternatives exist and can compensate in some tumors. Here, using genetically-engineered lung cancer model, we show that ablation of G6PD significantly suppresses *Kras^G12D/+^;Lkb1^-/-^*(KL) but not *Kras^G12D/+^;p53^-/-^* (KP) lung tumorigenesis. *In vivo* isotope tracing and metabolomics revealed that G6PD ablation significantly impaired NADPH generation, redox balance and *de novo* lipogenesis in KL but not KP lung tumors. Mechanistically, in KL tumors, G6PD ablation caused p53 activation that suppressed tumor growth. As tumor progressed, G6PD-deficient KL tumors increased an alternative NADPH source, serine-driven one carbon metabolism, rendering associated tumor-derived cell lines sensitive to serine/glycine depletion. Thus, oncogenic driver mutations determine lung cancer dependence on G6PD, whose targeting is a potential therapeutic strategy for tumors harboring KRAS and LKB1 co-mutations.

## Introduction

Tumor cells use nicotinamide adenine dinucleotide phosphate (NADPH) for redox homeostasis and reductive synthesis reactions to sustain their survival and growth ^1,2^. Consumption and production of NADPH are compartmentalized in the mitochondria and cytosol ^3,4^. Cytosolic NADPH is recycled through reduction of NADP+ via the oxidative pentose phosphate pathway (oxPPP) enzymes glucose 6-phosphate dehydrogenase (G6PD) and 6-phosphogluconate dehydrogenase (6PGD), isocitrate dehydrogenase 1 (IDH1), malic enzyme 1 (ME1), and the one-carbon metabolism (folate) enzymes methylenetetrahydrofolate dehydrogenase 1 (MTHFD1) and aldehyde dehydrogenase 1 family member L1 (ALDH1L1) ^3,5^. The functional importance of different metabolic enzymes involved in cytosolic NADPH homeostasis are not fully understood in cancer *in vivo*. Better understanding may open therapeutic opportunities.

Pentose phosphate pathway (PPP) flux is the major alternative glucose catabolic pathway to glycolysis. Dysregulation of proteins in this pathway is associated with cancer development, with the master antioxidant transcription factor NRF2 frequently upregulated in human cancer and driving oxPPP gene expression ^6,7^. The oxPPP pathway is essential for mammals, with knockout of the committed enzyme G6PD embryonic lethal. G6PD deficiency is the most common human enzyme defect because it protects against malaria ^8^. G6PD is upregulated in many cancers, and G6PD deficiency is associated with lower cancer risk and mortality for some cancers ^9–12^, suggesting that cancer cells may depend on G6PD for survival or proliferation. Loss of p53 upregulates G6PD activity and promotes NADPH-driven biosynthetic processes including *de novo* lipogenesis ^13^. In mouse models, G6PD deficiency significantly reduces melanoma metastasis^14^. Recently, we employed modern genetic tools to evaluate the role of G6PD in lung, breast, and colon cancer driven by oncogenic KRAS. We found that, in the studied KRAS mutant tumor models, G6PD, at most modestly promotes disease progression and is not strictly essential for solid tumorigenesis or metastatic spread ^15^. In particular, G6PD is not required for *Kras^G12D/+^;p53^-/-^* (KP) lung tumorigenesis ^15^. However, tumors KP tumors further lacking KEAP1, a tumor suppressor whose loss elevates NRF2, show greater dependence on G6PD ^7^. Thus, G6PD is likely to be particularly important in the context of specific tumor types or driver mutations.

Oncogenic KRAS mutations in non-small cell lung cancer (NSCLC) patients confer a poor prognosis and a high risk of cancer recurrence. LKB1 signaling negatively regulates tumor growth through direct phosphorylation and activation of the central metabolic sensor, AMPK, which governs glucose and lipid metabolism in response to alterations in nutrients and intracellular energy levels ^16–19^. Loss of LKB1 reprograms cancer cell metabolism to efficiently generate energy and biomass components for uncontrolled proliferation and dissemination. Meanwhile, such alterations in turn cause tumor cells to have less plasticity in response to metabolic stress, creating a metabolic vulnerability ^20,21^. TP53 and LKB1 co-mutations represent two different subgroups of KRAS-driven NSCLC, with distinct biological properties, metabolic vulnerabilities, and responses to standard therapies ^22–25^. Here, we explored whether the dispensability of G6PD for tumorigenesis is a common feature across different subtypes of KRAS-driven NSCLC. Compared to KP lung cancers, G6PD shows greater functional importance in *Kras^G12D/+^;Lkb1^-/-^* (KL) lung cancers. We observed that G6PD-deficient KL lung tumors showed increased oxidative stress, inducing p53 activation-mediated apoptosis and cell cycle arrest. G6PD loss also impaired *de novo* lipogenesis, whereas fat supplementation rescued the growth of G6PD-deficient KL lung tumors. Moreover, G6PD loss reprograms the NADPH generating metabolic pathway by increasing serine uptake to sustain one carbon metabolism-mediated cytosolic NADPH generation. G6PD-deficient KL lung tumor-derived cell lines (TDCLs) are sensitive to serine/glycine depletion, which is associated with increased reactive oxygen species (ROS). Thus, the dependence of G6PD-mediated oxPPP on KRAS-driven lung tumorigenesis is determined by specific oncogenic driver mutations.

## Results

### G6PD expression level correlates with the survival of lung cancer patients carrying KRAS and LKB1 co-mutations

Tumors exhibit an enormous demand for NADPH due to uncontrolled proliferation. The generation of cytosolic NADPH is primarily by metabolic enzymes such as G6PD, IDH1, ME1, and MTHFD1. Review of the cBioPortal datasets gave us further insight to conduct an overall survival comparison between low and high expression levels of cytosolic NADPH generating enzymes in lung cancer patients with KRAS wild type (WT) and KRAS mutations (MUT). In lung cancer patients with KRAS WT, except for MTHFD1, the mRNA expression levels of G6PD, IDH1, and ME1 were not correlated with the patient survival time (Fig. 1a). In patients with KRAS MUT, high mRNA expression levels of G6PD and MTHFD1 were associated with poorer overall survival (Fig. 1b). We further analyzed these correlations in patients carrying KRAS/TP53 co-mutations (co-MUT) and KRAS/LKB1 co-MUT. We found that high mRNA expression levels of G6PD and MTHFD1 were associated with poorer overall survival in patients with KRAS/LKB1 co-MUT (Fig. 1c), but not in patients with KRAS/TP53 co-MUT (Fig. 1d). In addition, G6PD mRNA expression is significantly higher in KRAS/LKB1 co-MUT lung cancer compared to KRAS/TP53 co-MUT lung cancer (Fig. 1e). These results suggest that G6PD and MTHFD1 expression impact survival in a subset of lung cancer patients (KRAS/LKB1 co-MUT and not KRAS/TP53 co-MUT lung cancers).

**Figure 1.**
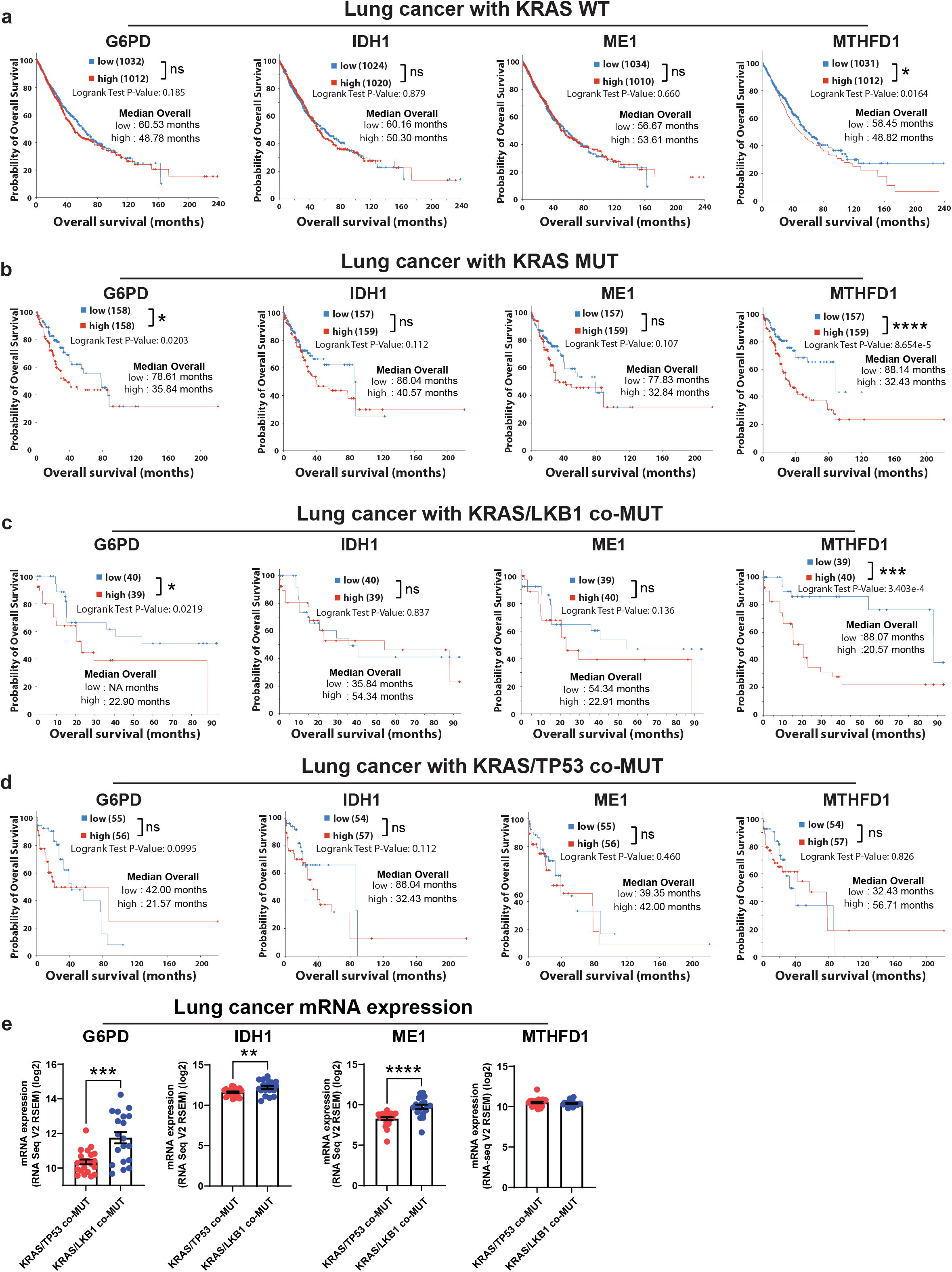
High G6PD mRNA expression correlates with poor survival of lung cancer patients with KRAS/LKB1 co-mutations. a. Overall survival comparison between low expression and high expression group of G6PD, IDH1, ME1 and MTHFD1 mRNA in lung cancer patients with KRAS wild type (WT), using data obtained from cBioPortal datasets. b. Overall survival comparison between low expression and high expression group of G6PD, IDH1, ME1 and MTHFD1 mRNA in lung cancer patients with KRAS mutations (MUT), using data obtained from cBioPortal datasets. c. Overall survival comparison between low expression and high expression group of G6PD, IDH1, ME1 and MTHFD1 mRNA in lung cancer patients with KRAS/LKB1 co-mutations (co-MUT), using data obtained from cBioPortal datasets. d. Overall survival comparison between low expression and high expression group of G6PD, IDH1, ME1 and MTHFD1 mRNA in lung cancer patients with KRAS/TP53 co-MUT, using data obtained from cBioPortal datasets. e. The mRNA expression levels of G6PD, ME1, IDH1, and MTHFD1 in lung cancer patients with KRAS/TP53 co-MUT and KRAS/LKB1 co-MUT, using data obtained from cBioPortal datasets. * *P*<0.05; ** *P*<0.01; *** *P*<0.005; **** *P*<0.001, ns: not significant

### G6PD is not essential for KP lung tumorigenesis

We previously found that conditional deletion of G6PD in KP lung tumor via CRISPR/Cas9-mediated gene editing *in vivo* did not affect KP lung tumorigenesis ^15^. We further validated this observation by generating a *G6pd^flox/flox^;Kras^LSL-G12D/+^;p53^flox/flox^*(*G6pd^flox/flox^;KP*) genetically engineered mouse model (GEMM) and concurrently deleting *G6pd* and inducing lung tumor via intranasal delivery of Lenti-Cre (Fig. 2a). G6PD expression was completely knocked out in KP lung tumors, which was validated by immunohistology (IHC) (Fig. 2b). We confirmed that G6PD is not essential for KP lung tumorigenesis by comparing the gross lung pathology (Fig. 2c), wet lung weight (Fig. 2d), quantification of tumor number (Fig. 2f) and tumor burden (Fig. 2g) of scanned lung hematoxylin & eosin (H&E) staining (Fig. 2e), tumor cell proliferation (Ki67) (Fig. 2h) and mouse survival (Fig. 2i) between mice bearing *G6pd^WT^;KP* lung tumors versus *G6pd^KO^;KP* lung tumors. Subsequently, we assessed the pool size levels of NADPH, NADP+, and the ratio of NADPH/NADP+, revealing comparable between *G6pd^WT^;KP* and *G6pd^KO^;KP* lung tumors (Fig. 2j). Consistent with these findings, G6PD loss in KP lung tumors also didn’t result in significant alternations in the levels of glutathione (GSH), glutathione disulfide (GSSG), and the ratio of GSH/GSSG (Fig. 2k). Taken together, we demonstrated that G6PD is dispensable for KP lung tumorigenesis and G6PD deficiency does not impact redox homeostasis in KP lung tumors.

**Figure 2.**
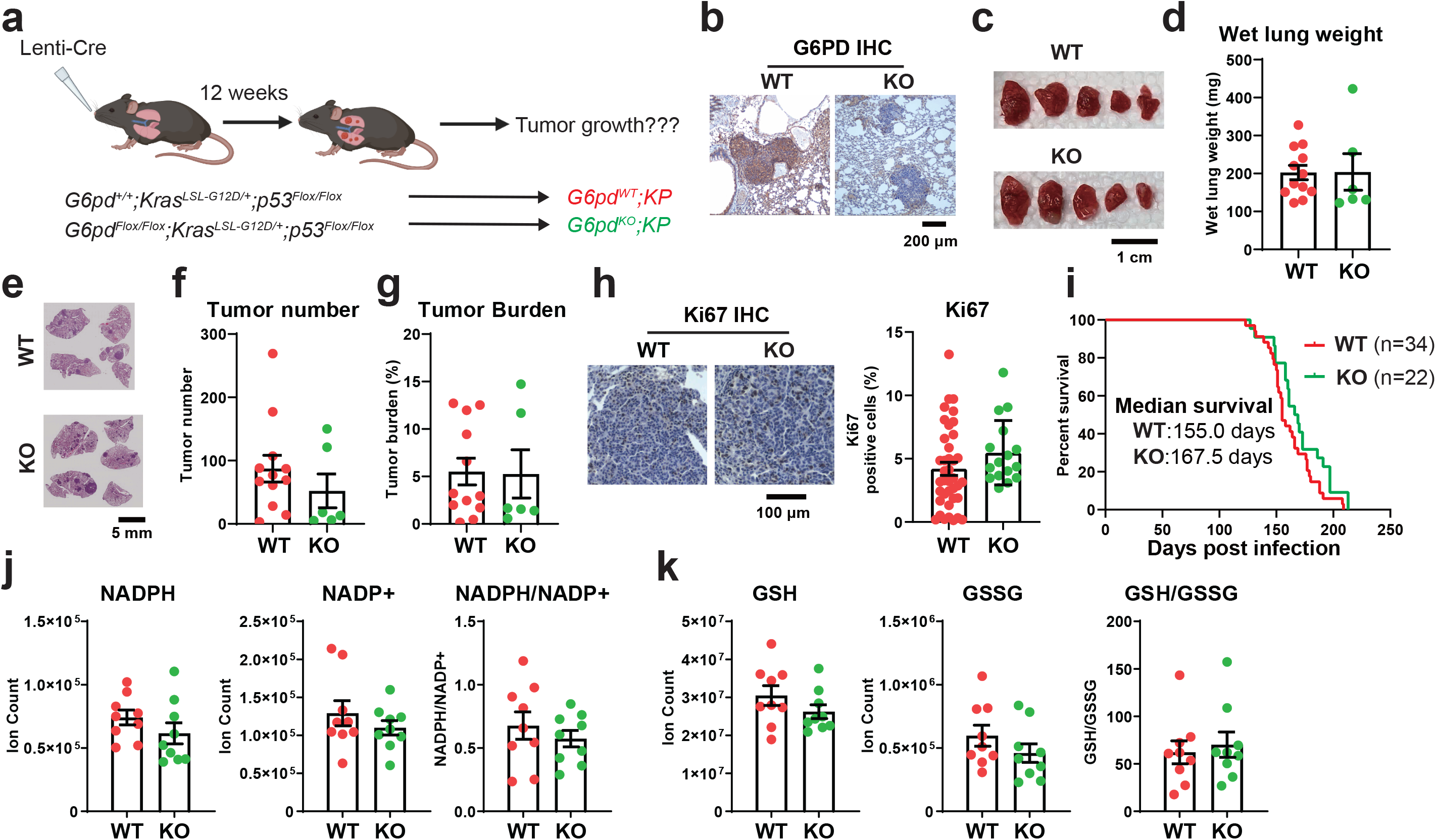
G6PD is not essential for KP lung tumorigenesis. a. Scheme to induce conditional tumoral *G6pd* knockout to study the role of G6PD in KP lung tumorigenesis. b. Representative immunohistochemistry (IHC) images of G6PD in *G6pd^WT^;KP* and *G6pd^KO^;KP* lung tumors. c. Representative gross lung pathology from mice bearing *G6pd^WT^;KP* and *G6pd^KO^;KP* lung tumors at 12 weeks post-tumor induction. d. Graph of wet lung weight from mice bearing *G6pd^WT^;KP* and *G6pd^KO^;KP* lung tumors at 12 weeks post-tumor induction. e. Representative histology (H&E staining) of scanned lung sections from (c). f & g. Quantification of tumor number (f) and tumor burden (g) from (e). h. Representative IHC images and quantification of Ki67 in *G6pd^WT^;KP* and *G6pd^KO^;KP* lung tumors. i. Kaplan-Meier survival curve of mice bearing *G6pd^WT^;KP* or *G6pd^KO^;KP* lung tumors. *P* > 0.05, log-rank test. j. Pool size of NADPH and NADP+, and NADPH/NADP+ ratio in *G6pd^WT^;KP* and *G6pd^KO^;KP* lung tumors at 12 weeks post-tumor induction. k. Pool size of glutathione (GSH) and glutathione disulfide (GSSG), and GSH/GSSG ratio in *G6pd^WT^;KP* and *G6pd^KO^;KP* lung tumors at 12 weeks post-tumor induction. Data are mean± SEM. For D_2_O infusion, n=3 mice for each group.

### G6PD promotes KL lung tumorigenesis

Co-MUT of TP53 and LKB1 represent two different subgroups of KRAS-driven lung cancer ^22–25^. To determine whether G6PD is dispensable for LKB1 mutation KRAS-driven (KL) NSCLC, as observed in KP lung tumors, we generated *G6pd^flox/flox^;Kras^LSL-G12D/+^;Lkb1^flox/flox^* (*G6pd^flox/flox^;KL*) GEMM of KL NSCLC and examined the role of G6PD in KL lung tumorigenesis (Fig. 3a). G6PD deletion in KL lung tumors was confirmed by IHC (Fig. 3b). We then performed a time course study of KL lung tumorigenesis at 7 and 12 weeks post-tumor induction. G6PD loss significantly reduced tumor number and tumor burden at 7 weeks, which was sustained at 12 weeks post-tumor induction (Fig. 3c-g). Consistent with these phenotypes, reduced cell proliferation (Ki67) (Fig. 3h) and reduced RAS downstream signaling (pErk and pS6) were observed in *G6pd^KO^;KL* lung tumors compared to *G6pd^WT^;KL* tumors (Fig. 3i). Moreover, mice bearing *G6pd^KO^;KL* lung tumors had a significantly longer life span compared to mice bearing *G6pd^WT^;KL* lung tumors (Fig. 3j). In summary, we demonstrated that in contrast to KP lung tumors G6PD supports KL lung tumor initiation and growth.

**Figure 3.**
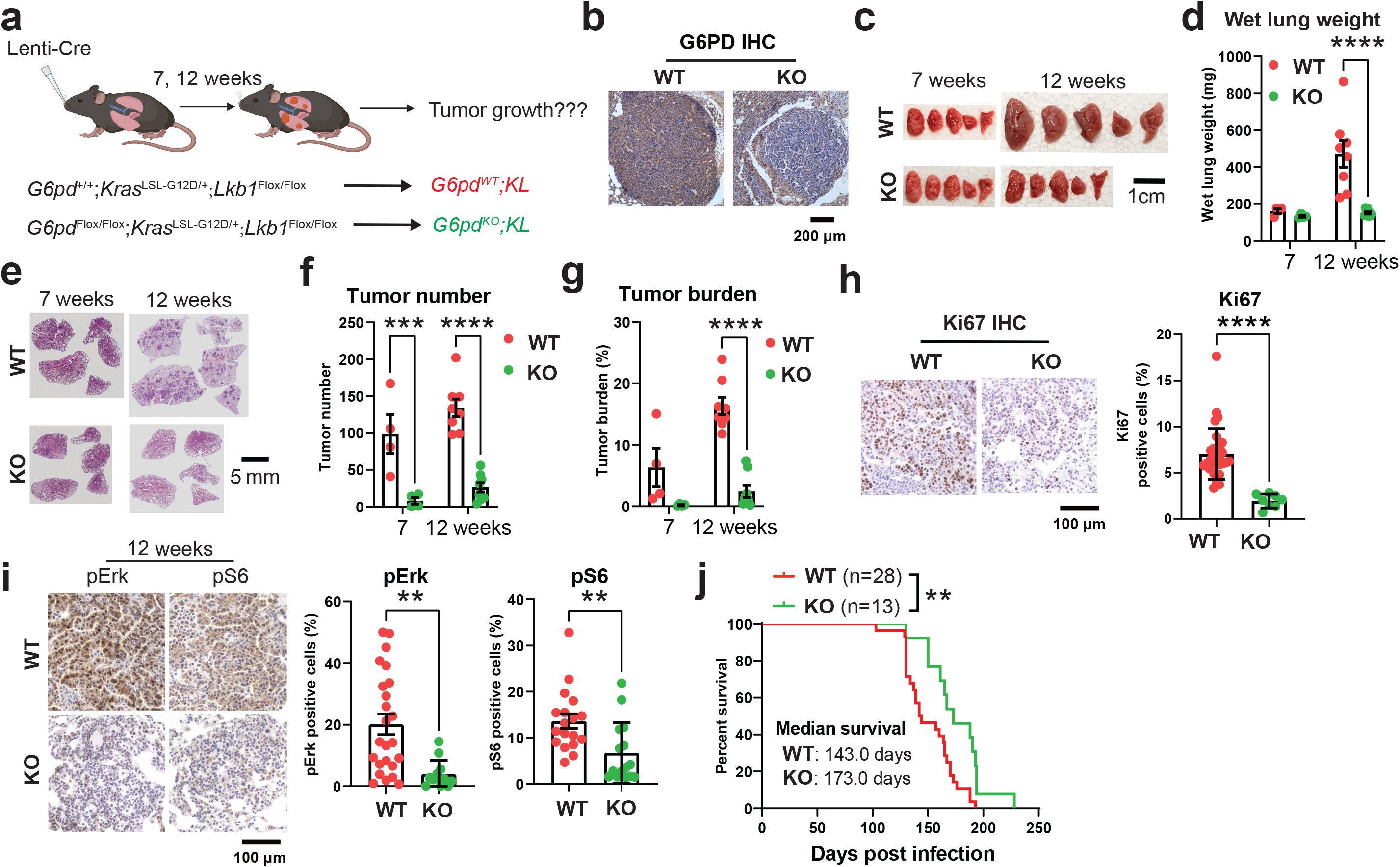
G6PD promotes KL lung tumorigenesis. a. Scheme to induce conditional tumoral *G6pd* knockout to study the role G6PD in KL lung tumorigenesis. b. Representative IHC images of G6PD in *G6pd^WT^;KL* and *G6pd^KO^;KL* lung tumors at 12 weeks post-tumor induction. c. Representative gross lung pathology from mice bearing *G6pd^WT^;KL* and *G6pd^KO^;KL* lung tumors at 7 and 12 weeks post-tumor induction. d. Graph of wet lung weight from (c). e. Representative histology (H&E staining) of scanned lung sections from (c). f & g. Quantification of tumor number (f) and tumor burden (g) from (e). h. Representative IHC images and quantification of Ki67 in *G6pd^WT^;KL* and *G6pd^KO^;KL* lung tumors at 12 weeks post-tumor induction. i. Representative IHC images and quantification of pErk, and pS6 in *G6pd^WT^;KL* and *G6pd^KO^;KL* lung tumors at 12 weeks post-tumor induction. j. Kaplan-Meier survival curve of mice bearing *G6pd^WT^;KL* or *G6pd^KO^;KL* lung tumors. *P*<0.01, log-rank test. Data are mean± SEM. * *P*<0.05; ** *P*<0.01; *** *P*<0.005; **** *P*<0.001.

### G6PD is involved in generating NADPH in KL lung tumors, thereby maintaining cellular redox homeostasis

G6PD-mediated oxPPP is one of the cytosolic NADPH generating metabolic pathways. We hypothesized that G6PD depletion would suppress NADPH production in KL lung tumors, disrupting redox homeostasis and leading to cell death in the stressed tumor microenvironment. As expected, the level of NADPH in *G6pd^KO^;KL* lung tumors was significantly lower than *G6pd^WT^;KL* lung tumors, leading to an impaired redox balance in *G6pd^KO^;KL* lung tumors, evidenced by a lower ratio of NADPH/NADP+ and GSH/GSSG in *G6pd^KO^;KL* lung tumors than *G6pd^WT^;KL* (Fig. 4a, b). Moreover, Gene Set Enrichment Analysis (GSEA) of bulk-tumor mRNA-seq revealed that oxidative stress signaling was significantly upregulated in *G6pd^KO^;KL* lung tumors in contrast to *G6pd^WT^;KL* lung tumors (Fig. 4c). This was further confirmed by IHC of 8-oxo-dG and γ-H2AX, markers for oxidative stress. Both 8-oxo-dG and γ-H2AX were significantly increased in *G6pd^KO^;KL* lung tumors compared to *G6pd^WT^;KL* lung tumors (Fig. 4d).

**Figure 4.**
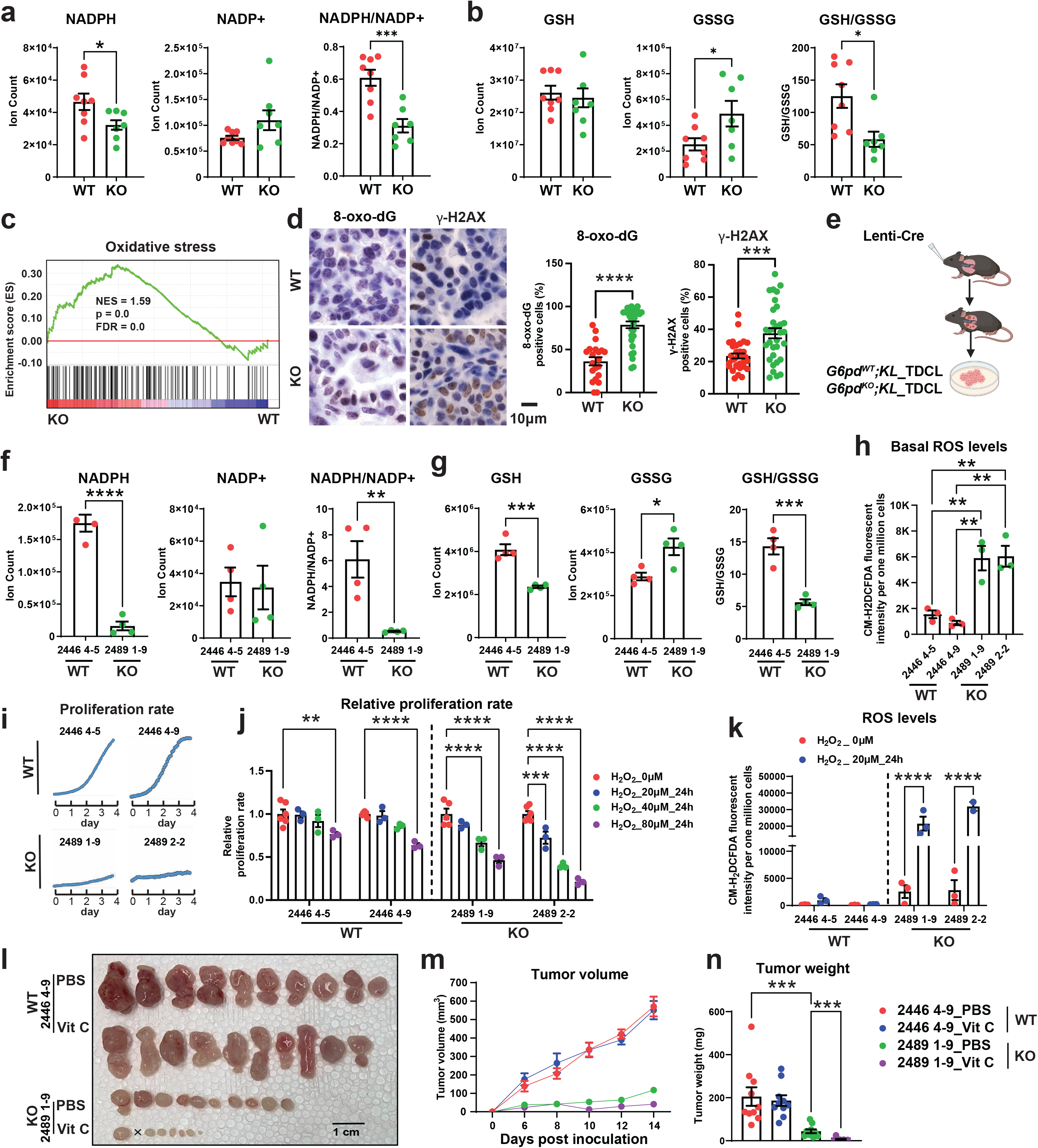
G6PD-mediated NADPH production is essential to maintain cellular redox homeostasis in KL tumorigenesis. a. Pool size of NADPH and NADP+, and NADPH/NADP+ ratio in *G6pd^WT^;KL* and *G6pd^KO^;KL* lung tumors at 12 weeks post-tumor induction. b. Pool size of GSH and GSSG, and GSH/GSSG ratio in *G6pd^WT^;KL* and *G6pd^KO^;KL* lung tumors at 12 weeks post-tumor induction. c. Gene Set Enrichment Analysis (GSEA) of oxidative stress signaling for *G6pd^WT^;KL* and *G6pd^KO^;KL* lung tumors based on bulk-tumor mRNA-seq data. d. Representative IHC images and quantification of 8-oxo-dG and γ-H2AX in *G6pd^WT^;KL* and *G6pd^KO^;KL* lung tumors at 12 weeks post-tumor induction. e. Scheme of generating tumor-derived cell lines (TDCLs) from *G6pd^WT^;KL* and *G6pd^KO^;KL* lung tumors. f. Pool size of NADPH and NADP+, and NADPH/NADP+ ratio of *G6pd^WT^;KL* and *G6pd^KO^;KL* TDCLs in nutrient rich conditions (complete RPMI medium). g. Pool size of GSH and GSSG, and GSH/GSSG ratio of *G6pd^WT^;KL* and *G6pd^KO^;KL* TDCLs in nutrient rich conditions. h. Basal ROS level of *G6pd^WT^;KL* and *G6pd^KO^;KL* TDCLs in nutrient rich conditions. i. Proliferation rate of *G6pd^WT^;KL* and *G6pd^KO^;KL* TDCLs in nutrient rich conditions for 4 days. j. Relative proliferation rate of *G6pd^WT^;KL* and *G6pd^KO^;KL* TDCLs treated with different concentration of H_2_O_2_ for 24 hours. k. ROS levels of *G6pd^WT^;KL* and *G6pd^KO^;KL* TDCLs treated with 20 μmol/L H_2_O_2_ for 24 hours. l. Allograft tumor growth curve of *G6pd^WT^;KL* and *G6pd^KO^;KL* TDCLs treated with or without high-dose Vitamin C (Vit C, 4g/kg, *i.p*, daily) for 2 weeks. m. Gross pathology of *G6pd^WT^;KL* and *G6pd^KO^;KL* allograft tumors at the end of the experiment from (l). n. *G6pd^WT^;KL* and *G6pd^KO^;KL* allograft tumor weight at the end of the experiment from (l). Data are mean± SEM. * *P*<0.05; ** *P*<0.01; *** *P*<0.005; **** *P*<0.001.

To further deduce the consequences of damaged redox homeostasis on KL lung tumor growth, we generated *G6pd^WT^;KL* and *G6pd^KO^;KL* TDCLs from mouse lung tumors (Fig. 4e). In nutrient-rich conditions, *G6pd^KO^;KL* TDCLs displayed lower NADPH, lower ratio of NADPH/NADP+ and GSH/GSSG, and higher basal ROS levels than *G6pd^WT^;KL* TDCLs (Fig. 4f-h). We further observed that the proliferation rate of *G6pd^KO^;KL* was significantly lower than *G6pd^WT^;KL* TDCLs (Fig. 4i). Moreover, compared with *G6pd^WT^;KL* TDCLs, *G6pd^KO^;KL* TDCLs were more sensitive to H_2_O_2_-induced cell death (Fig. 4j), which was associated with a significant increase in ROS levels (Fig. 4k). Finally, we performed *in vivo* TDCLs-induced allograft tumor growth assays to determine whether *G6pd^KO^;KL* allograft tumors is more sensitive to oxidative stress than *G6pd^WT^;KL* allograft tumors, as we observed in *in vitro* cell culture. High-doses of Vitamin C (Vit C) has been reported to induce oxidative stress in preclinical mouse models and has been proposed in clinical studies combined with standard therapies ^26^. Therefore, we treated mice bearing *G6pd^WT^;KL* or *G6pd^KO^;KL* allograft tumors with vehicle control or high-dose Vit C (4g/kg). As expected, G6PD deficiency significantly reduced allograft tumor growth in the vehicle control group. Importantly, a high dose of Vit C further suppressed the growth of KL allograft tumors only if they also lacked G6PD (Fig. 4l-n). In summary, we demonstrated that G6PD-mediated NADPH production maintains KL lung tumor redox homeostasis to prevent oxidative stress-induced cell death and support tumor growth.

### G6PD loss activates p53 to suppress KL lung tumorigenesis

DNA damage and oxidative stress activate p53, leading to cell cycle arrest and apoptosis ^27^. GSEA of bulk-tumor mRNA-seq showed that the p53-mediated apoptotic pathway was upregulated in *G6pd^KO^;KL* lung tumors compared to *G6pd^WT^;KL* tumors (Fig. 5a). This was further validated by IHC of p53 and its downstream targets, p21 and cleaved caspase-3, which were significantly upregulated in *G6pd^KO^;KL* lung tumors compared to *G6pd^WT^;KL* tumors (Fig. 5b). We subsequently proposed that loss of G6PD in KL lung tumors induces p53 activation, leading to growth arrest and apoptosis. To test this hypothesis, we generated *G6pd^flox/flox^;Kras^LSL-G12D/+^;p53^flox/flox^;Lkb1^flox/flox^*(*G6pd^flox/flox^;KPL*) and *G6pd^+/+^;KPL* mice, and examined the effect of G6PD ablation on KPL lung tumorigenesis (Fig. 5c). G6PD deficiency in KPL lung tumors was confirmed by IHC (Fig. 5d). We found that p53 loss eliminated the sensitivity of KL lung tumors to G6PD knockout. This was measured by gross lung pathology, wet lung weight, quantification of tumor number and tumor burden from scanned lung H&E sections at six weeks post KPL lung tumor induction (Fig. 5e-i). Moreover, tumor cell proliferation (Ki67) was comparable between *G6pd^WT^;KPL* and *G6pd^KO^;KPL* lung tumors (Fig. 5j). As a result, the life spans of mice bearing *G6pd^WT^;KPL* and *G6pd^KO^;KPL* lung tumors were similar (Fig. 5k). Thus, the slow growth of G6PD-knockout KL tumors is due to oxidative stress-inducing p53 activation and p53 activation inhibiting tumor progression.

**Figure 5.**
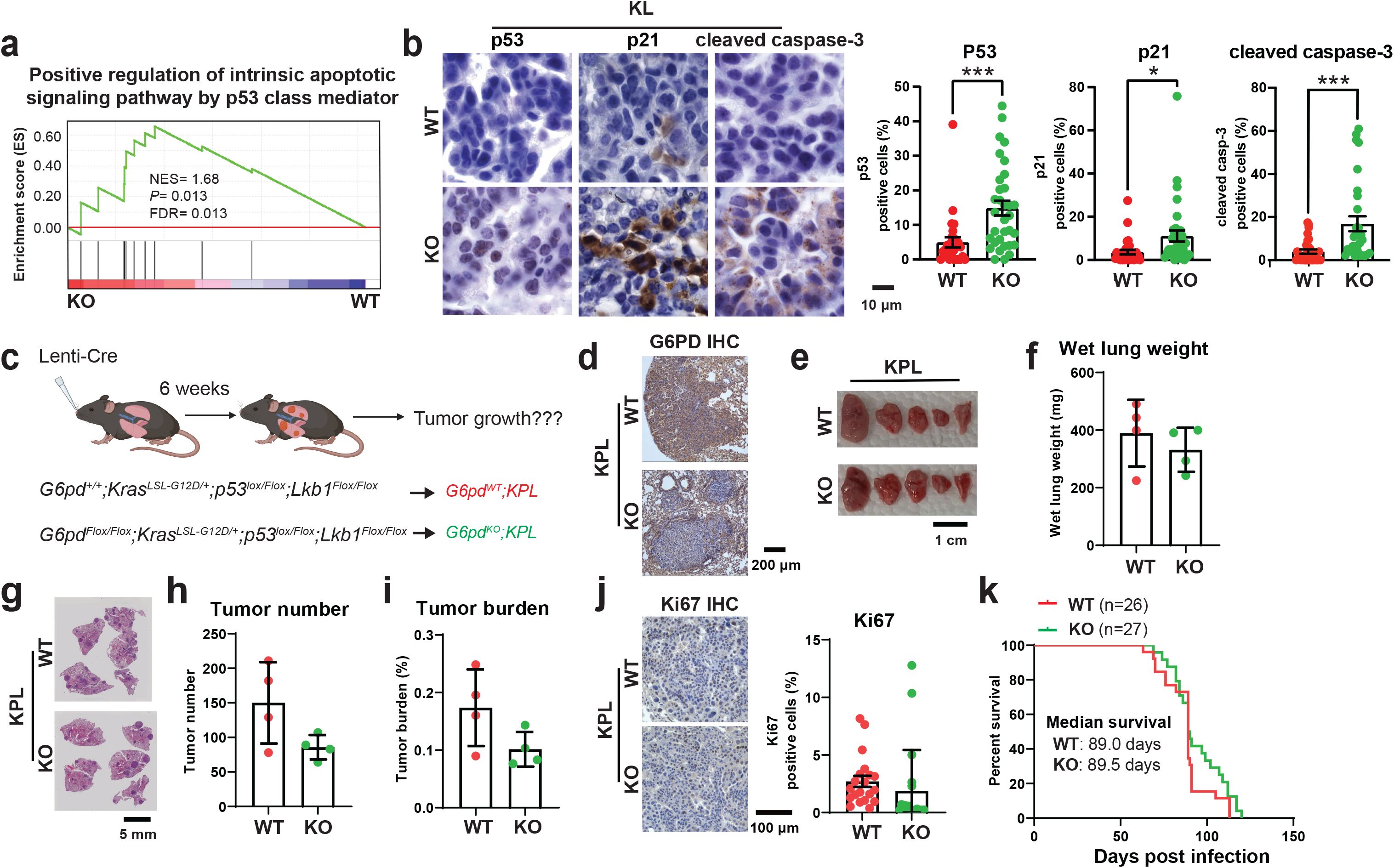
G6PD suppresses p53 activation for KL lung tumorigenesis. a. GSEA of positive regulation of intrinsic apoptotic signaling pathway by p53 class mediator for *G6pd^WT^;KL* and *G6pd^KO^;KL* lung tumors based on bulk-tumor mRNA-seq data. b. Representative IHC images and quantification of p53, p21 and cleaved caspase-3 of *G6pd^WT^;KL* and *G6pd^KO^;KL* lung tumors at 12 weeks post-tumor induction. c. Scheme to induce conditional tumoral *G6pd* knockout to study the role of G6PD in KPL lung tumorigenesis. d. Representative IHC images of G6PD in *G6pd^WT^;KPL* and *G6pd^KO^;KPL* lung tumors at 6 weeks post-tumor induction. e. Representative gross lung pathology from mice bearing *G6pd^WT^;KPL* and *G6pd^KO^;KPL* lung tumors at 6 weeks post-tumor induction. f. Graph of wet lung weight from mice bearing *G6pd^WT^;KPL* and *G6pd^KO^;KPL* lung tumors at 6 weeks post-tumor induction. g. Representative histology (H&E staining) of scanned lung sections from mice bearing *G6pd^WT^;KPL* and *G6pd^KO^;KPL* lung tumors at 6 weeks post-tumor induction. h & i. Quantification of tumor number (h) and tumor burden (i) from (g). j. Representative IHC images and quantification of Ki67 in *G6pd^WT^;KPL* and *G6pd^KO^;KPL* lung tumors. k. Kaplan-Meier survival curve of mice bearing *G6pd^WT^;KPL* and *G6pd^KO^;KPL* lung tumors. *P*>0.05, log-rank test. Data are mean± SEM. * *P*<0.05; *** *P*<0.005.

### G6PD depletion impairs KL lung tumor lipid metabolism

Cytosolic NADPH provides a hydride source for *de novo* fatty acid synthesis. Pathway analysis of bulk-tumor mRNA-seq of KL lung tumors revealed that G6PD ablation significantly altered lipid metabolism by downregulating the pathways related to lipid and fatty acid biosynthesis (Fig. 6a, b). We further performed lipidomics of lung tumors and serum from tumor-bearing mice. In order to investigate the fatty acyl composition in the lipids, we extracted and saponified the lipids to release the fatty acyl chains for the LC-MS analysis. We found that G6PD loss altered the fatty acyl composition in KL lung tumors when in a fasted state (Fig. 6c, d), but not in a fed state (Supplemental Fig. 1a, b). Compared with *G6pd^WT^;KL* lung tumors, *G6pd^KO^;KL* tumors had significantly lower levels of long-chain fatty acyl groups, whereas very long-chain fatty acyl groups accumulated in *G6pd^KO^;KL* lung tumors (Fig. 6d). G6PD loss in KP lung tumors did not impact the fatty acyl composition (Supplemental Fig. 1c, d). Moreover, there was no difference in the fatty acyl composition in serum between mice bearing *G6pd^WT^;KL* and *G6pd^KO^;KL* lung tumors (Fig. 6c, d).

**Figure 6.**
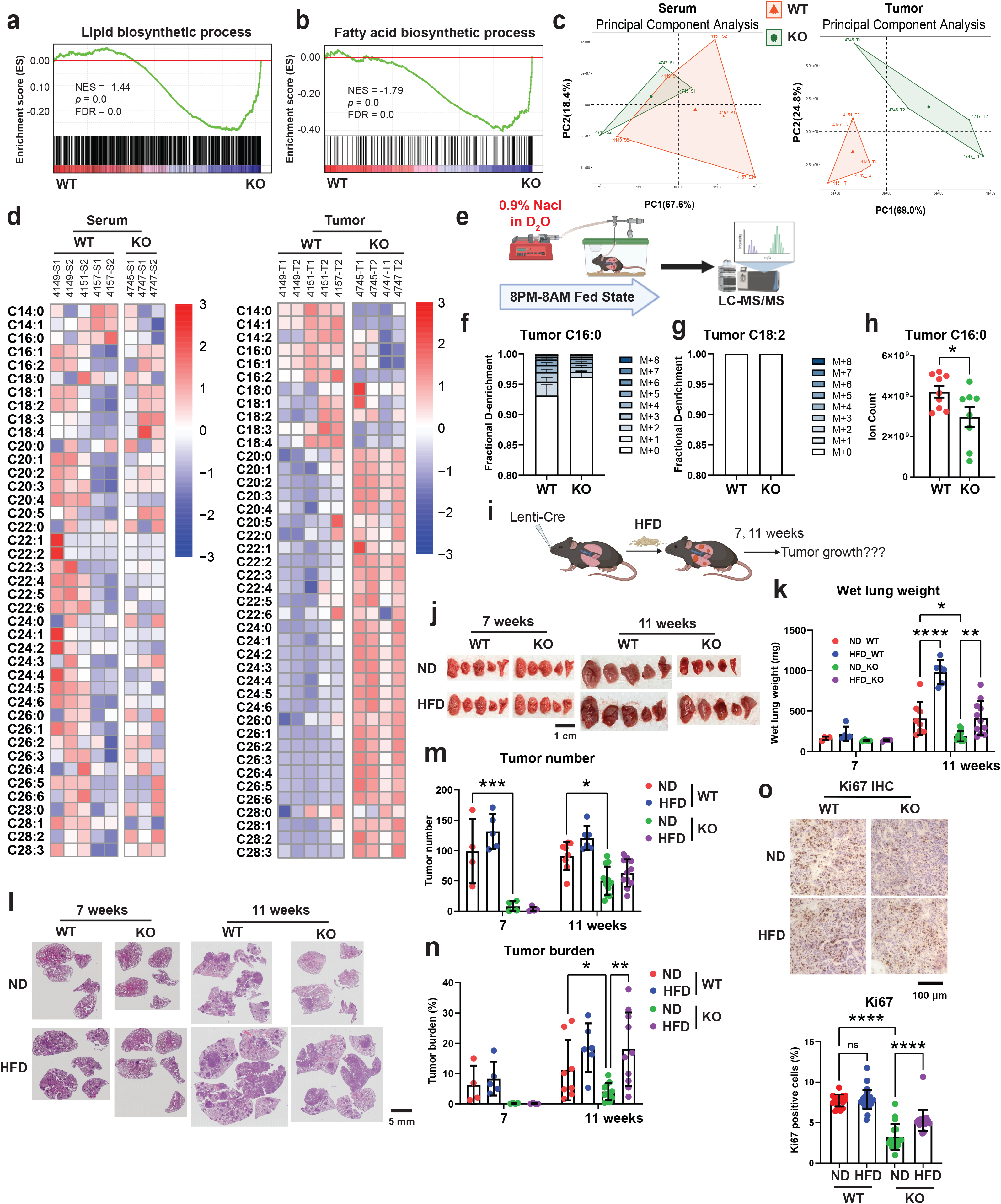
G6PD depletion impairs KL lung tumor lipid metabolism. a & b. GSEA of lipid biosynthetic process (a) and fatty acids biosynthetic process (b) for *G6pd^WT^;KL* and *G6pd^KO^;KL* lung tumors based on bulk-tumor mRNA-seq data. c & d. Principal Component Analysis (PCA) (c) and Heatmap (d) of saponified fatty acids pool size of *G6pd^WT^;KL* and *G6pd^KO^;KL* lung tumors and serum from KL lung tumor bearing mice at fasted state (food was removed from the mice at approximately 9:00 a.m., and mice were euthanized and tumor samples were collected at 3:00 p.m.) at 12 weeks post-tumor induction. e. Scheme of *in vivo* D_2_O infusion to examine KL lung tumor *de novo* fatty acid synthesis. f & g. C16:0 (f) and C18:2 (g) deuterium (^2^H) labeling fraction in *G6pd^WT^;KL* and *G6pd^KO^;KL* lung tumors after 12 hour D_2_O infusion at 12 weeks post-tumor induction. h. C16:0 pool size of *G6pd^WT^;KL* and *G6pd^KO^;KL* lung tumors at 12 weeks post-tumor induction. i. Scheme to examine the impact of high-fat diet (HFD) on *G6pd^WT^;KL* and *G6pd^KO^;KL* lung tumorigenesis. j. Representative gross lung pathology from mice bearing *G6pd^WT^;KL* and *G6pd^KO^;KL* lung tumors fed with normal diet (ND) or HFD at 7, 11 weeks post-tumor induction. k. Graph of wet lung weight from mice bearing *G6pd^WT^;KL* and *G6pd^KO^;KL* lung tumors fed with ND or HFD at 7, 11 weeks post-tumor induction. l. Representative histology (H&E staining) of scanned lung sections from mice bearing *G6pd^WT^;KL* and *G6pd^KO^;KL* lung tumors fed with ND or HFD at 7, 11 weeks post-tumor induction. m & n. Quantification of tumor number (m) and tumor burden (n) from (l). o. Representative IHC images and quantification of Ki67 in *G6pd^WT^;KL* or *G6pd^KO^;KL* lung tumors at 11 weeks post-tumor induction. Data are mean± SEM. * *P*<0.05; ** *P*<0.01; **** *P*<0.0001.

D_2_O has long been used as a tracer for assessing *in vivo* lipogenesis ^28–30^, because ^2^H can be incorporated onto fatty acids primarily via deuterated NADPH exchanged with ambient D_2_O ^31^. We subsequently examined the *de novo* fatty acid synthesis of KL lung tumors by intravenously infusing D_2_O into mice bearing *G6pd^WT^;KL* or *G6pd^KO^;KL* lung tumors via jugular vein for 12 hours (8:00 p.m. - 8:00 a.m., fed state) (Fig. 6e) and assessed the fatty acid labeling from D_2_O. Lower ^2^H labeling in C16:0 in *G6pd^KO^;KL* lung tumors than in *G6pd^WT^;KL* tumors was observed (Fig. 6f). As expected, no ^2^H labeling was detected in essential fatty acid C18:2 (Fig. 6g). Moreover, the level of C16:0 was significantly lower in *G6pd^KO^;KL* lung tumors than in *G6pd^WT^;KL* tumors (Fig.6h). Glucose provides carbon building blocks for *de novo* fatty acid synthesis. Using [U-^13^C_6_]-glucose as a tracer, we observed significantly lower *de novo* fatty acid synthesis in *G6pd^KO^;KL* TDCLs than in *G6pd^WT^;KL* TDCLs (Supplemental Fig. 2a, b), which is consistent with ^2^H-labeled *de novo* lipogenesis *in vivo* (Fig. 6f).

Subsequently, we tested the hypothesis that lowering NADPH impairs *de novo* lipogenesis, which may lead to less availability of fatty acids for *G6pd^KO^;KL* tumor growth by supplementing mice with high-fat diet (HFD) and examining tumor burden at 11 weeks post-tumor induction (Fig. 6i). HFD successfully rescued the KL lung tumor growth and cell proliferation (Ki67) caused by G6PD depletion (Fig. 6j-o). However, HFD did not alter tumor number in mice bearing *G6pd^KO^;KL* lung tumors (Fig. 6m), indicating that lower fat availability due to reduced *de novo* lipogenesis may not affect KL lung tumor initiation, but suppress tumor growth. Thus, during KL lung tumorigenesis, oxPPP-mediated NADPH production is essential for maintaining lipid metabolism for KL lung tumor growth.

### G6PD ablation does not affect TCA cycle metabolism but reprograms serine metabolism

Glucose contributes carbon to TCA cycle metabolites, PPP, and non-essential amino acids ^6^. G6PD oxidizes the glycolytic intermediate G6P to 6-phosphogluconolactone. Therefore, we performed *in vivo* [U-^13^C6]-glucose tracing and metabolic flux analysis in mice bearing *G6pd^WT^;KL* or *G6pd^KO^;KL* lung tumors to determine whether G6PD ablation in KL lung tumors has any impact on glucose metabolism besides oxPPP (Fig. 7a). We found that glucose carbon flux to KL lung tumor glycolytic intermediates, and TCA cycle metabolites and derivatives was not affected by G6PD ablation (Fig. 7b, c). The levels of G6P and 3-phosphoglycerate (3-PG) were significantly higher in *G6pd^KO^;KL* lung tumors than *G6pd^WT^;KL* tumors. However, the levels of pyruvate, lactate and TCA cycle metabolites were comparable between *G6pd^WT^;KL* and *G6pd^KO^;KL* lung tumors (Fig. 7d, e).

**Figure 7.**
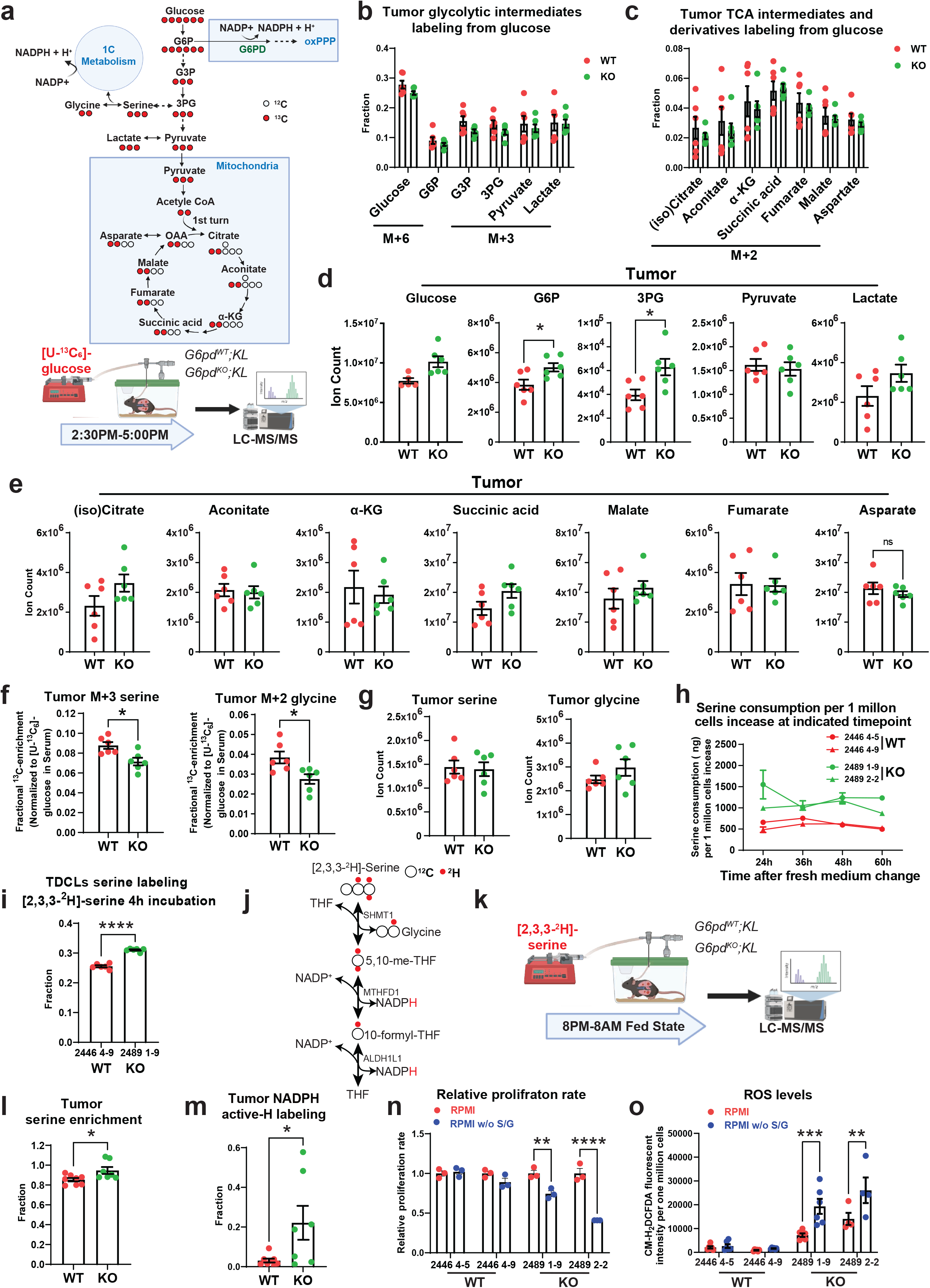
G6PD ablation has no impact on KL TCA cycle metabolism, but reprograms serine metabolism. a. Scheme of carbon contribution from glucose to glycolytic intermediates, TCA cycle intermediates, and serine (top) and *in vivo* [U-^13^C_6_]-glucose tracing in the mice bearing *G6pd^WT^;KL* and *G6pd^KO^;KL* lung tumors (bottom). b. Normalized ^13^C labeling fraction of glycolytic intermediates from glucose of *G6pd^WT^;KL* and *G6pd^KO^;KL* lung tumors in fasted state at 12 weeks post-tumor induction. c. Normalized ^13^C labeling fraction of TCA cycle intermediates from glucose of *G6pd^WT^;KL* and *G6pd^KO^;KL* lung tumors in fasted state at 12 weeks post-tumor induction. d. Pool size of glycolytic intermediates of *G6pd^WT^;KL* and *G6pd^KO^;KL* lung tumors in fasted state at 12 weeks post-tumor induction. e. Pool size of TCA cycle metabolites of *G6pd^WT^;KL* and *G6pd^KO^;KL* lung tumors in fasted state at 12 weeks post-tumor induction. f. Normalized ^13^C labeling fraction from glucose to serine and glycine of *G6pd^WT^;KL* and *G6pd^KO^;KL* lung tumors in fasted state at 12 weeks post-tumor induction. g. Pool size of serine and glycine of *G6pd^WT^;KL* and *G6pd^KO^;KL* lung tumors in fasted state at 12 weeks post-tumor induction. h. Serine consumption of *G6pd^WT^;KL* and *G6pd^KO^;KL* TDCLs in nutrient rich conditions. i. ^2^H labeling fraction of serine in *G6pd^WT^;KL* and *G6pd^KO^;KL* TDCLs after 4 hours [2,3,3-^2^H]-serine labeling in nutrient rich conditions. j. Scheme of hydrogen contribution from [2,3,3-^2^H]-serine to NADPH. k. Scheme of *in vivo* [2,3,3-^2^H]-serine tracing in the mice bearing *G6pd^WT^;KL* and *G6pd^KO^;KL* lung tumors. l. ^2^H labeling fraction of serine in *G6pd^WT^;KL* and *G6pd^KO^;KL* lung tumors at 12 weeks post-tumor induction. m. NADPH active-H labeling from [2,3,3]-serine in *G6pd^WT^;KL* and *G6pd^KO^;KL* lung tumors at 12 weeks post-tumor induction. n. Relative proliferation rate of *G6pd^WT^;KL* and *G6pd^KO^;KL* TDCLs cultured with RPMI media with or without serine and glycine for 48 hours. o. ROS levels of *G6pd^WT^;KL* and *G6pd^KO^;KL* TDCLs cultured with RPMI media with or without serine and glycine for 48 hours. Data are mean± SEM. * *P*<0.05; ** *P*<0.01; *** *P*<0.005; **** *P*<0.001.

Glucose also contributes carbons for biosynthesis by incorporating glycolytic intermediates into different metabolic pathways. The synthesis of serine from glucose is a key metabolic pathway supporting cellular proliferation in healthy and malignant cells. We found that the ^13^C labeling of serine and glycine from glucose was significantly lower in *G6pd^KO^;KL* lung tumors than in *G6pd^WT^;KL* tumors, suggesting that G6PD ablation impairs serine biosynthesis (Fig. 7f). The reduction of glucose carbon flux to serine could contribute to the higher 3-PG levels in *G6pd^KO^;KL* lung tumors than in *G6pd^WT^;KL* tumors (Fig. 7d). However, the overall levels of serine and glycine in KL lung tumors were not altered by G6PD ablation (Fig. 7g). This could be compensated by the upregulation of uptake or the reduction of catabolism. To distinguish these two possibilities, we first examined the serine consumption of KL TDCLs by measuring the reduction of serine in conditioned medium. We found that serine consumption in *G6pd^KO^;KL* TDCLs was significantly higher than *G6pd^WT^;KL* TDCLs (Fig. 7h). Subsequently, we assessed serine utilization via *in vitro* isotope tracing using [2,2,3-^2^H]-serine and observed a higher serine ^2^H labeling in *G6pd^KO^;KL* TDCLs than *G6pd^WT^;KL* TDCLs (Fig. 7i), further demonstrating that G6PD deficiency reprograms KL tumor cell metabolism by upregulating serine uptake.

Next, we performed [2,2,3- ^2^H]-serine *in vivo* tracing in KL lung tumor-bearing mice (Fig. 7j, k) and observed significantly higher serine enrichment in *G6pd^KO^;KL* lung tumors than *G6pd^WT^;KL* lung tumors (Fig. 7l), confirming an increased serine uptake *in vivo* due to G6PD loss. Moreover, compared to *G6pd^WT^;KL* lung tumors, a significantly increased ^2^H labeling onto NADPH in *G6pd^KO^;KL* tumors was observed (Fig. 7m).

Finally, we cultured *G6pd^WT^;KL* and *G6pd^KO^;K*L TDCLs in serine/glycine-free RPMI medium to determine the consequence of reprogrammed serine metabolism on cell proliferation. Compared to *G6pd^WT^;KL* TDCLs, *G6pd^KO^;KL* TDCLs were much more sensitive to serine/glycine depletion-induced cell death (Fig. 7n). Moreover, serine/glycine depletion significantly increased ROS levels in *G6pd^KO^;KL* TDCLs, but not in *G6pd^WT^;KL* TDCLs (Fig. 7o). Our data suggests that a reprogramed serine metabolism by G6PD loss could be used for NADPH generation, thereby maintaining redox homeostasis.

## Discussion

Cellular pools of NADP(H) are compartmentalized ^2^. *In vitro* studies have demonstrated that cytosolic and mitochondrial NADPH fluxes are independently and precisely regulated by multiple metabolic pathways ^5,32^. G6PD-mediated oxPPP is one of the metabolic pathways involved in cytosolic NADPH generation. In this study, by using GEMMs of KRAS-driven NSCLC, we found that the dependence on G6PD is distinct in different subtypes of lung cancer. G6PD promotes KL but not KP lung tumorigenesis. LKB1 serves as a central modifier of cellular response to different metabolic stress. Loss of LKB1-AMPK signaling results in heightened sensitivity to energy depletion and to disturbances in redox homeostasis ^33^. It is possible that KL lung tumors, which lack proper AMPK activity, exhibit a greater metabolic vulnerability and less plasticity in response to G6PD loss when compared to KP lung tumors that retain intact LKB1 function.

The roles of G6PD in cancer development depend on its metabolic function in producing NADPH to reduce ROS and to support reductive biosynthesis ^1,2^. *De novo* fatty acid synthesis is needed for growth and viability of NSCLC cells. Energy regulator AMPK inhibits the activity of Acetyl-CoA carboxylase (ACC) to suppress *de novo* fatty acid synthesis and promote fatty acid oxidation ^33^. An allosteric inhibitor of the ACC enzymes ACC1 and ACC2 markedly suppressed KL lung tumor growth ^34^. We observed that G6PD deficiency has a substantial impact on NADPH levels in KL lung tumors. This disruption impairs redox homeostasis and reduces *de novo* fatty acid synthesis. As a consequence, G6PD-deficient KL lung tumor cells are more sensitive to oxidative stress compared to WT cells. Moreover, high-fat diet supplementation rescued KL lung tumor growth caused by G6PD ablation, indicating that less fat availability due to reduced *de novo* fatty acid synthesis may contribute to the slower growth of *G6pd^KO^;KL* lung tumors. Our study further demonstrates the role of G6PD-mediated oxPPP in cancer, which is specific to KL but not KP NSCLC.

Cancer cells exhibit aberrant redox homeostasis, but while ROS are pro-tumorigenic, high ROS levels are cytotoxic ^27^ and can be induced by oncogenic activity ^35^. p53 influences the cellular redox balance by regulating several genes with antioxidant or pro-oxidant properties, which depends on various factors, including p53 protein levels ^36^. Moderately elevated ROS levels inhibit p53, while higher levels promote its expression ^36^. Active p53 and its downstream targets p21 and cleaved caspase-3 are significantly upregulated in *G6pd^KO^;KL* lung tumors compared to *G6pd^WT^;KL* tumors. This could be due to G6PD loss-induced oxidative stress and high ROS levels. Therefore, during KL lung tumorigenesis, G6PD-mediated NADPH production modulates oxidative stress to suppress p53 activation, thereby supporting tumor growth.

Cancer cells frequently alter metabolism to adapt to challenges. Inhibition of one cytosolic NADPH-producing metabolic pathway may lead to upregulation of others to compensate. Indeed, *in vitro* cell culture studies show that cancer cells can tolerate the loss of any two of the four canonical cytosolic NADPH production routes ^37^. More studies are needed to explore compensatory mechanisms between NADPH-generating routes to maintain cytosolic NADPH homeostasis for KP lung tumorigenesis *in vivo*. As for KL lung tumors, G6PD ablation did not completely inhibit the growth of KL lung tumors, suggesting the presence of metabolic reprogramming or compensation. Our *in vivo* isotope tracing and flux analysis revealed that G6PD deficiency in KL lung tumors does not affect glucose carbon flux to tumor pyruvate, lactate, and TCA cycle intermediates. However, G6PD loss in KL lung tumors reduces glucose carbon flux to serine. Additionally, serine uptake is increased to maintain the serine level in G6PD-deficient KL lung tumors for cytosolic NADPH production. We found that in *in vitro* cell culture, increased serine uptake is used to maintain redox homeostasis for cell proliferation. This is different from KEAP1 mutant tumors, in which G6PD loss triggered TCA intermediate depletion because of up-regulation of the alternative NADPH-producing enzymes malic enzyme and isocitrate dehydrogenase ^7^.

G6PD has been proposed as a potential therapeutic target for cancer therapy in recent years due to its overexpression in various cancers ^38^. G6PD inhibitors have been sought to achieve this goal ^2,38,39^. Based on the data from the cBioPortal datasets, our analysis suggests high G6PD expression is associated with the poor survival of lung cancer patients with co-MUT of KRAS and LKB1. Therefore, patients harboring KRAS and LKB1 co-MUT may benefit from G6PD inhibitor therapy. Moreover, the G6PD inhibitor combined with the reagents that induce oxidative stress can be a potential therapeutic strategy to treat this subtype of KRAS-mutant NSCLC.

## Materials and Methods

### cBioPortal data processing

The overall survival analysis comparing the low and high expression levels of cytosolic NADPH generating enzymes, including G6PD, IDH1, ME1, and MTHFD1, in lung cancer patients was conducted using the cBioPortal datasets (https://www.cbioportal.org/, accessed on June 25, 2023) ^40^. Data from 33 studies (as listed in Table 1) available in the cBioPortal datasets was utilized for the present analysis.

**Table 1.** List of the 33 studies involved in the overall survival analysis for lung cancer in the cBioPortal datasets.

| No. | study |
| --- | --- |
| 1 | Lung Adenocarcinoma (Broad, Cell 2012) |
| 2 | Lung Adenocarcinoma (CPTAC, Cell 2020) |
| 3 | Lung Adenocarcinoma (MSK Mind,Nature Cancer 2022) |
| 4 | Lung Adenocarcinoma (MSK, 2021) |
| 5 | Lung Adenocarcinoma (MSK, J Thorac Oncol 2020) |
| 6 | Lung Adenocarcinoma (MSK, NPJ Precision Oncology 2021) |
| 7 | Lung Adenocarcinoma (MSK, Science 2015) |
| 8 | Lung Adenocarcinoma (OncoSG, Nat Genet 2020) |
| 9 | Lung Adenocarcinoma (TCGA, Firehose Legacy) |
| 10 | Lung Adenocarcinoma (TCGA, Nature 2014) |
| 11 | Lung Adenocarcinoma (TCGA, PanCancer Atlas) |
| 12 | Lung Adenocarcinoma (TSP, Nature 2008) |
| 13 | Lung Adenocarcinoma Met Organotropism (MSK, Cancer cell 2023) |
| 14 | Lung Cancer (SMC, Cancer Research 2016) |
| 15 | Lung Cancer in Never Smokers (NCI, Nature Genetics 2021) |
| 16 | Lung Squamous Cell Carcinoma (CPTAC, Cell 2021) |
| 17 | Lung Squamous Cell Carcinoma (TCGA, Firehose Legacy) |
| 18 | Lung Squamous Cell Carcinoma (TCGA, Nature 2012) |
| 19 | Lung Squamous Cell Carcinoma (TCGA, PanCancer Atlas) |
| 20 | Metastatic Non-Small Cell Lung Cancer (MSK, Nature Medicine 2022) |
| 21 | Non-Small Cell Cancer (MSK, Cancer Discov 2017) |
| 22 | Non-Small Cell Lung Cancer (MSK, Cancer Cell 2018) |
| 23 | Non-Small Cell Lung Cancer (MSK, J Clin Oncol 2018) |
| 24 | Non-Small Cell Lung Cancer (MSK, Science 2015) |
| 25 | Non-Small Cell Lung Cancer (TRACERx, NEJM & Nature 2017) |
| 26 | Non-Small Cell Lung Cancer (University of Turin, Lung Cancer 2017) |
| 27 | Pan-Lung Cancer (TCGA, Nat Genet 2016) |
| 28 | Small Cell Lung Cancer (CLCGP, Nat Genet 2012) |
| 29 | Small Cell Lung Cancer (Johns Hopkins, Nat Genet 2012) |
| 30 | Small Cell Lung Cancer (U Cologne, Nature 2015) |
| 31 | Small-Cell Lung Cancer (Multi-Institute, Cancer Cell 2017) |
| 32 | Thoracic Cancer (MSK, Nat Commun 2021) |
| 33 | Thoracic PDX (MSK, Provisional) |

For the overall survival analysis, the “gene specific” option was chosen, adding gene names including G6PD, IDH1, ME1, and MTHFD1. The mRNA data type selected was “mRNA expression z-scores relative to all samples (log RNA Seq V2 RSEM)“. A chart was then generated to compare the two groups based on the median expression of the indicated gene’s mRNA. Subsequently, the overall survival was examined between the mRNA low expression group and the mRNA high expression group of the indicated gene.

For the analysis of mRNA expression levels of G6PD, IDH1, ME1, and MTHFD1, data were obtained from a 586 samples study on lung cancer (Lung Adenocarcinoma, TCGA, Firehose Legacy) available in the cBioPortal datasets. Sample information for those with KRAS/TP53 co-MUT and KRAS/LKB1 co-MUT was extracted from the study, and the mRNA expression levels of the indicated genes were compared between these two groups. The mRNA expression levels were represented as mRNA expression z-scores relative to all samples (log RNA Seq V2 RSEM).

### Mice

Rutgers Animal Care and Use Committee (IACUC) has approved all animal experiments performed in this study. *Kras^G12D^;Lkb1^flox/flox^*mice, *Kras^G12D^;p53^flox/flox^* mice generated in our previous study ^41,42^, and *G6pd^flox/flox^*mice generated by Rutgers Cancer Institute of New Jersey Genome Editing core facility, underwent cross-breeding with each other to generate *G6pd^flox/flox^;Kras^LSL-G12D^;Lkb1^flox/flox^ (G6pd^flox/flox^;KL), _G6pdflox/flox;KrasLSL-G12D;p53flox/flox (G6pdflox/flox;KP)_*_, *Kras*_*_LSL-G12D;p53flox/flox;Lkb1flox/flox (KPL)_*_, and *G6pd*_*_flox/flox;KrasLSL-G12D;Lkb1flox/flox;p53flox/flox (G6pdflox/flox;KPL)_* _mice._

At 6-8 weeks of age, mice were intranasally infected with Lenti-Cre (University of Iowa Viral Vector Core) at 5×10^6^ plaque-forming units (pfu) per mouse to induce lung tumors, following the methodology employed in our previous investigation ^41^.

### Histology and IHC

Mice were euthanized via cervical dislocation at the designated time points following Lenti-Cre infection. Lung tissues were collected and placed in formaldehyde (Fisher Scientific, #SF93-4) for a period of 12-24 hours. Afterward, the tissues were transferred to a 70% ethanol solution and stored at 4°C. Paraffin-embedded sections were prepared using the methodology described in a previous study for H&E staining and IHC ^43^. The antibodies employed for IHC were G6PD (Abcam, #AB993), Ki67 (Abcam, #ab15580), pS6 (Cell Signaling, #4858S), P-p42/44 MAPK (p-ERK) (Cell Signaling, #9101S), cleaved caspase3 (Cell Signaling, #9661S), p53 (Leica, #DO-7), p21 (Santa Cruz Biotech, #sc-6246), 8-oxo-dG (R&D systems, #4354-MC-050), γ-H2AX (Cell Signaling, #9718).

For quantification of IHC for Ki67, pS6, p-ERK, cleaved caspase3, p53, p21, 8-oxo-dG, γ-H2AX, 10-50 representative images from each group were obtained and scored using the ImageJ software.

### Tumor number/burden quantification

H&E-stained lung specimens were imaged using an Olympus VS120 whole-slide scanner (Olympus Corporation of the Americas) at 20 × magnification at the Rutgers Cancer Institute of New Jersey Biomedical Informatics shared resource. Image analysis was conducted using a custom-developed protocol on the Visiopharm image analysis platform (Visiopharm A/S). The protocol facilitated the identification of tissue area and the computation of tumor burden based on semiautomatically detected tumors. Low-resolution image maps, extracted from the whole-slide images, were utilized to generate tumor masks and whole-tissue masks. These masks were generated for each slide, enabling the segmentation of tumor burden ratios.

### High fat diet treatment

*G6pd^WT^;KL* and *G6pd^KO^;KL* lung tumors were induced by intranasal infection with Lenti-Cre. On the same day of infection, half of mice were fed with the high-fat diet (Bio-Serv Mouse Diet, #F3282) for a duration of 11 weeks and half were fed with the laboratory control diet (Bio-Serv Mouse Diet, #S4207). After the 11 weeks treatment period, the mice were euthanized, and lung tissues were collected for H&E staining, tumor number/burden quantification and IHC.

### D_2_O, [U-^13^C_6_]-glucose and [2,2,3-^2^H]-serine infusion

Before the infusion experiments, venous catheters were surgically implanted into the jugular veins of tumor-bearing mice, with a 3 to 4 days interval. The infusions were conducted on conscious, freely moving mice. For the infusion of D_2_O (Cambridge Isotope, #DLM-4-50) and [2,2,3-^2^H]-serine (Cambridge Isotope, #DLM-582-0.5), mice were fed continuously throughout the infusion period (8:00 p.m. - 8:00 a.m.). For the infusion of [U-^13^C_6_]-glucose (Cambridge Isotope, #CLM-1396-1), food was removed from the mice at approximately 9:00 a.m., and infusion was commenced at 3:00 p.m.. Mice were infused with D_2_O saline (0.9% NaCl) at a rate of 0.1 mL/g/min or [2,2,3-^2^H]-serine (200 mmol/L) at a rate of 0.2 mL/g/min for 12 hours overnight, or [U-^13^C_6_]-glucose (200 mmol/L) at a rate of 0.1 mL/g/min for 2.5 hours before being euthanized for rapid lung tumors collection. Blood samples for serum analysis were collected from the mice’s cheeks into 1.5 mL Eppendorf Tubes (Flex-Tubes, #20901-551). Lung tumors were swiftly dissected and frozen using a liquid-nitrogen cold clamp to halt metabolic activity and then stored at -80°C until further metabolites extraction.

### mRNA-seq and GSEA analysis

*G6pd^WT^;KL* and *G6pd^KO^;KL* lung tumors were induced by intranasal infection with Lenti-Cre. At 12 weeks post-tumor induction, mice were euthanized by cervical dislocation. The lung tumors were rapidly dissected and snap-frozen in liquid nitrogen. Subsequently, the frozen samples were pulverized to a powder using a Cryomill (Retsch). High-quality total RNA was extracted from the above samples, and mRNA enrichment were performed using RNeasy Min Kit (QIAGEN, #74104). cDNA library was prepared and sequenced at Novogene.

For GSEA analysis, gene sets for “Oxidative stress,” “Positive regulation of intrinsic apoptotic signaling pathway by p53 class mediator“, “Lipid biosynthetic process“, and “Fatty acid biosynthetic process” were downloaded from the MSigDB website (https://www.gsea-msigdb.org/). A dataset containing mRNA expression profiles of all genes for *G6pd^WT^;KL* and *G6pd^KO^;KL* lung tumors was prepared. The GSEA v4.3.2 software, using the classic setting recommended for mRNA-seq data in the GSEA manual, was employed to perform the GSEA analysis.

### TDCLs and cell culture

*G6pd^WT^;KL* or *G6pd^KO^;KL* TDCLs were generated from *G6pd^WT^;KL* or *G6pd^KO^;KL* lung tumors at 12 weeks post-tumor induction. TDCLs were cultured in complete RPMI medium (Gibco, #11875-093) supplemented with 10% fetal bovine serum (FBS), 1% Penicillin-Streptomycin, and 0.075% sodium bicarbonate) at 37°C with 5% CO_2_. Regular testing using the Universal mycoplasma detection kit (ATCC, #30-1012k) confirmed the absence of mycoplasma contamination in the cell lines.

### Allograft mouse model and high-dose Vit C treatment

For allograft tumor induction, *G6pd^WT^;KL* or *G6pd^KO^;KL* lung TDCLs were subcutaneously injected into the right and left flank of male C57BL/6 mice at 1 × 10^6^ cells/injection at 6-8 weeks of age.

For the induction of an increase in ROS in allograft tumors, mice bearing allograft tumors were administered Vit C at a dosage of 4 g/kg intraperitoneally (*i.p.*) daily for 2 weeks. Tumor volume was measured using a caliper every other day during the 2-week treatment period. After the two weeks of treatment, the mice were euthanized, and tumors were collected and weighed for further analysis.

### Cell proliferation assay

For IncuCyte measurement, cells were seeded at a density of 4×10^4^ cells per well in 12-well plates under a complete RPMI medium. Cell proliferation assays were conducted over 4 days using IncuCyte live-cell analysis system, and the data were analyzed using IncuCyte Zoom System.

For manual cell counting, cells were treated with H_2_O_2_ (Sigma-Aldrich, #88597-100ML-F) at concentrations of 0, 20, 40, and 80 μmol/L for 24 hours. Subsequently, the cells were trypsinized off the culture plates and counted using a Vi-cell XR cell viability analyzer (Beckman coulter). The relative proliferation rate for cells treated with different concentrations of H_2_O_2_ was calculated by normalizing the cell number to the corresponding cells without H_2_O_2_ treatment.

### Serine consumption assay

*G6pd^WT^;KL* or *G6pd^KO^;KL* TDCLs were seeded at 0.5 × 10^5^ or 1 × 10^5^ cells per well in a 24-well plate in complete RPMI medium, respectively. The following day, fresh complete RPMI medium was replaced, and medium was collected at 0, 24, 36, 48, and 60 hours. Each timepoint set up duplicate wells for both *G6pd^WT^;KL* and *G6pd^KO^;KL* TDCLs. The levels of serine in medium were measured using LC-MS. A Vi-cell XR cell viability analyzer was used to measure cell number at each time point. Based on the following formula: the reduction in serine amount in the well (serine amount at the 0-hour timepoint minus the serine amount at the indicated timepoint) divided by the increase in cell number in the same well (cell number at the indicated timepoint minus the cell number at the 0-hour timepoint), the serine consumption (μg) per one million cells increase at the indicated timepoint can be calculated.

### Serine and glycine depletion assay

*G6pd^WT^;KL* and *G6pd^KO^;KL* TDCLs were cultured in complete RPMI medium (RPMI medium without glucose, serine, and glycine (Teknova, #R9660) supplemented with 10% fetal FBS, 1% Penicillin-Streptomycin, 0.075% sodium bicarbonate, 2g/L glucose (Sigma, #G8270-1KG), 10 mg/L glycine (Sigma, #50046-50G) and 30 mg/L serine (Sigma, #S4311-25G)) at 37°C with 5% CO_2_. After 2 days, *G6pd^WT^;KL* TDCLs were trypsinized and seeded at 0.5 × 10^5^ cells per well, while *G6pd^KO^;KL* TDCLs were trypsinized and seeded at 1 × 10^5^ cells per well in a 24-well plate. For Serine and glycine depletion assay, the TDCLs were cultured in the serine/glycine free RPMI medium (RPMI medium without glucose, serine, and glycine supplemented with 10% fetal FBS, 1% Penicillin-Streptomycin, 0.075% sodium bicarbonate, 2g/L glucose), and the complete RPMI medium as control. After 2 days, TDLCs were trypsinized off plates and then counted using a Vi-cell XR cell viability analyzer.

### ROS levels measurement

The CM-H_2_DCFDA assay (Invitrogen, #C6827) was performed to measure cellular ROS levels. *G6pd^WT^;KL* or *G6pd^KO^;KL* TDCLs were seeded at 0.5 × 10^5^ or 1 × 10^5^ cells per well in a 24-well plate in complete RPMI medium, respectively. After 2 days, the cells were washed twice with HBSS (Corning, #21-022-CV). Then, 0.5 mL of CM-H_2_DCFDA solution with a concentration of 5 μmol/L in HBSS was added to each well, and the cells were incubated at 37°C for 45 minutes in the dark. Following incubation, the medium was changed to RPMI medium (Gibco, #11875-093) for 30 minutes to allow for recovery. Subsequently, the medium was replaced with HBSS, and the fluorescence intensity was measured using a microplate reader (Tecan). The excitation wavelength was set to 493 nm, and the emission wavelength was set to 520 nm. After measuring the fluorescence intensity, TDLCs were trypsinized off the plates and counted using a Vi-cell XR cell viability analyzer. The ROS levels were calculated using the following formula: the fluorescence intensity (the fluorescence intensity of each well stained with CM-H_2_DCFDA minus the fluorescence intensity without staining) was divided by the cell number (x 10^6^) in the same well.

For the ROS levels measurement under the H_2_O_2_ treatment condition, cells were treated with H_2_O_2_ at 0, 20 μmol/L concentrations for 24 hours. Subsequently, the cells were stained with CM-H_2_DCFDA following the aforementioned method.

To measure ROS levels under serine and glycine depletion conditions, cells were treated according to the “serine and glycine depletion assay” method. Subsequently, the cells were stained with CM-H_2_DCFDA following the aforementioned method.

### Serine uptake measurement *in vitro*

*G6pd^WT^;KL* and *G6pd^KO^;KL* lung TDCLs were seeded in 6-cm dishes with regular complete RPMI medium for serine uptake measurement. The following day, the medium was replaced with the RPMI medium without glucose, serine, and glycine (Teknova, #R9660) supplemented with 10% FBS, 1% penicillin-streptomycin, 0.075% sodium bicarbonate, 10 mg/L glycine, and 30 mg/L [2,2,3-^2^H]-serine. The cells were incubated for 4 hours, and the experiment was performed in triplicate. Subsequently, water-soluble metabolites were extracted and subjected to LC-MS analysis to further analyze and calculate ^2^H labeling for serine as serine uptake.

### Sample preparation of water-soluble metabolites for LC-MS analysis

For the extraction of water-soluble metabolites from lung tumors, following the methodology described in a previous study ^44^. Approximately 20 mg of tumor samples were precisely weighed and placed into a pre-cooled tube. The tissue samples were then pulverized using the Cryomill. Pre-cooled extraction buffer consisting of methanol: acetonitrile: H_2_O (40:40:20, V/V) with 0.5% formic acid (Sigma-Aldrich, # F0507-100ML) was added to the resulting powder (40 μL of solvent per mg of tumors). The samples were then vortexed for 15 seconds and incubated on ice for 10 minutes. Subsequently, 15% NH_4_HCO_3_ solution (5% V/V of the extraction buffer) was used to neutralize the samples. Then all samples were vortexed again for 10 seconds and centrifuged at 4°C, 13,000 × g for 20 minutes. The resulting supernatant was transferred to LC-MS vials for subsequent analysis.

For the extraction of water-soluble metabolites from serum, following the methodology described in a previous study ^44^, pre-cooled methanol was added to the serum samples in a 1.5 mL Eppendorf Tube (Tube A). The mixture was vortexed for 10 seconds and left at -20°C for 20 minutes. Afterward, the tube was centrifuged at 4°C for 10 minutes. The top supernatant was carefully transferred to another 1.5 mL Eppendorf Tube (Tube B) as the first extract. Then, 200 μL of an extraction buffer composed of methanol: acetonitrile: H_2_O (40:40:20, V/V) was added to Tube A, which still contained the pellet at the bottom. The tube was vortexed for 10 s and placed on ice for 10 minutes. Subsequently, the tube was centrifuged at 4°C, 13,000 × g for 10 minutes. The top supernatant from this step was combined with the first extract in Tube B, resulting in a final extract volume of 250 μL. The extract was loaded onto the Phree Phospholipid Removal Tabbed 1mL tube (Phenomenex, #8B-S133-TAK) and centrifuged at 4°C, 13,000 × g for 10 minutes. The extract was transferred to LC-MS vials for subsequent analysis.

For the extraction of water-soluble metabolites from cultured cell samples, following the methodology described in a previous study ^41^, *G6pd^WT^;KL* and *G6pd^KO^;KL* TDCLs cultured in triplicate in 6-cm dishes were washed twice with PBS. Subsequently, the cells were incubated with 1 mL of pre-cooled extraction buffer containing methanol: acetonitrile: H_2_O in a ratio of 40:40:20, along with 0.5% formic acid solution, on ice for 5 minutes. The extraction process was followed by neutralization with 50 µL of 15% ammonium bicarbonate. The cells were then scraped from the plates and transferred to 1.5 mL tubes. Afterward, the tubes were centrifuged at 4°C, 13,000 × g for 10 minutes. The resulting supernatant was transferred to LC-MS vials for subsequent analysis.

### Sample preparation of saponified fatty acids for LC-MS analysis

To extract saponified fatty acids from lung tumors and serum, following the methodology described in a previous study ^45^, pre-cooled methanol was added to the resulting powder (12 μL of methanol per mg of lung tumors or μL of serum). The samples were vortexed for 10 seconds, followed by adding -20°C MTBE (40 μL of MTBE per mg of tumors or μL of serum). After another 10 seconds vortexing step, the samples were incubated on a shaker at 4°C for 6 min. Next, H_2_O (10 μL of H_2_O per mg of tumors or μL of serum) was added, and the samples were centrifuged at 4°C, 13,000 × g for 2 minutes. Following centrifugation, 160 μL of the top MTBE layer was transferred to a new 1.5 mL Eppendorf Tube (Flex-Tubes, #20901-551). The liquid was then dried under the air. Subsequently, the sample was resuspended in 1 mL of saponification solvent (0.3 mol/L KOH in 90:10 methanol/H_2_O), and the entire volume was transferred to 4 mL glass vials. The vials were placed in a water bath at 80°C for 1 hour. After incubation, the vials were cooled on ice for 3 minutes, and 100 μL of formic acid was added. Then, 300 μL of hexanes was added, and the samples were vortexed for 10 seconds, resulting in two layers. The top layer was transferred to a new 1.5 mL Eppendorf Tube, and this step was repeated to obtain a final volume of 600 μL. The liquid was then dried in the 1.5 mL Eppendorf Tube under air. To resuspend the extracted sample, 150 μL of resuspension solvent (50:50 isopropanol/methanol) was added. The samples were centrifuged at 4°C, 13,000 × g for 10 minutes, and the resulting supernatant was transferred to LC-MS vials for further analysis.

### LC-MS methods

For the LC-MS analysis of water-soluble metabolites, following the methodology described in a previous study ^46^, the experimental conditions were optimized using an HPLC-ESI-MS system consisting of a Thermo Scientific Vanquish HPLC coupled with a Thermo Q Exactive Plus MS. The HPLC system was equipped with a Waters XBridge BEH Amide column (2.1 mm × 150 mm, 2.5 μm particle size, 130Å pore size) and a Waters XBridge BEH XP VanGuard cartridge (2.1 mm × 5 mm, 2.5 μm particle size, 130Å pore size) guard column. The column temperature was maintained at 25°C. The mobile phase A consisted of a mixture of H_2_O: acetonitrile (95:5, V/V) with 20 mmol/L NH_3_AC and 20 mmol/L NH_3_OH at pH 9. The mobile phase B consisted of a mixture of acetonitrile: H_2_O (80:20, V/V) with 20 mmol/L NH_3_AC and 20 mmol/L NH_3_OH at pH 9. The composition of mobile phase B varied over time as follows: 0 min, 100%; 3 min, 100%; 3.2 min, 90%; 6.2 min, 90%; 6.5 min, 80%; 10.5 min, 80%; 10.7 min, 70%; 13.5 min, 70%; 13.7 min, 45%; 16 min, 45%; 16.5 min, 100%. The flow rate was set to 300 μL/min, and the injection volume was 5 μL. The column temperature was maintained at 25°C throughout the analysis. MS scans were acquired in negative ionization mode, with a resolution of 70,000 at m/z 200. The automatic gain control (AGC) target was set to 3 × 10^^6^, and the mass-to-charge ratio (m/z) scan range was set from 72 to 1000 ^45^.

For the LC-MS analysis of NADPH and NADP+, the gradient consisted of the following steps: 0 min, 85% B; 2 min, 85% B; 3 min, 60% B; 9 min, 60% B; 9.5 min, 35% B; 13 min, 5% B; 15.5 min, 5% B; 16 min, 85% B. The run was stopped at 20 mins, and the injection volume was 15 µL. As described previously, full scans were alternated with targeted scans in the m/z range of 640-765, with a resolution of 35,000 at m/z 200, and with AGC target of 5 × 10^5^.

For the LC-MS analysis of fatty acids samples, following the methodology described in a previous study ^41^, a Vanquish Horizon UHPLC system (Thermo Fisher Scientific, Waltham, MA) with a Poroshell 120 EC-C18 column (150 mm × 2.1 mm, 2.7 μm particle size, Agilent Infinity Lab, Santa Clara, CA) was employed using a gradient of solvent A (90%:10% H_2_O: methanol with 34.2 mmol/L acetic acid, 1 mmol/L ammonium acetate, pH 9.4), and solvent B (75%:25% IPA: methanol with 34.2 mmol/L acetic acid, 1 mmol/L ammonium acetate, pH 9.4). The gradient program was as follows: 0 min, 25% B; 2 min, 25% B; 5.5 min, 65% B; 12.5 min, 100% B; 19.5 min, 100% B; 20.0 min, 25% B; 30 min, 25% B. The flow rate was set to 200 μL/min, and the column temperature was maintained at 55°C. MS data was acquired using a Thermo Q Exactive PLUS mass spectrometer with heated electrospray ionization (HESI) source. The spray voltage was set to -2.7 kV in negative mode. The sheath gas, auxiliary gas, and sweep gas flow rates of 40, 10, and 2 (arbitrary unit), respectively. The capillary temperature was set to 300°C, and the auxiliary gas heater was set to 360°C. The S-lens RF level was 45. In negative ionization mode, the m/z scan range was set from 200 to 2,000. The AGC target was set to 1 × 10^^6^ and the maximum injection time (IT) was 200 ms. The resolution was set to 140,000. Data-dependent MS/MS scans were acquired from pooled samples in negative ionization mode with a loop count of 3, an AGC target of 1 × 10^^6^, and the maximum IT of 50ms. The mass resolution was set to 17,500, and the normalized collision energy (NCE) was stepped at 20, 30, and 40. Dynamic exclusion was set to 10.0 s. The MS/MS data were processed using MS-DIAL ^47,48^, and the fatty acids annotations were performed by matching the built-in MS/MS database.

### NADPH active-H labeling calculation

NADPH and NADP+ features were extracted in EI-MAVEN software. The ^2^H isotope natural abundance and impurity of labeled substrate were corrected using AccuCor written in R ^49^. NADPH active-H labeling ***p*** from [2,3,3-^2^H]-serine was determined based on labeling fraction of NADPH-NADP+ pair and calculated using the previously described ***equation (1)*** ^45^, the matrix on the left side of ***equation (1)*** contains the experimentally measured mass ^2^H isotope distribution for NADP+, while the right side contains the experimentally measured mass ^2^H isotope distribution for NADPH.

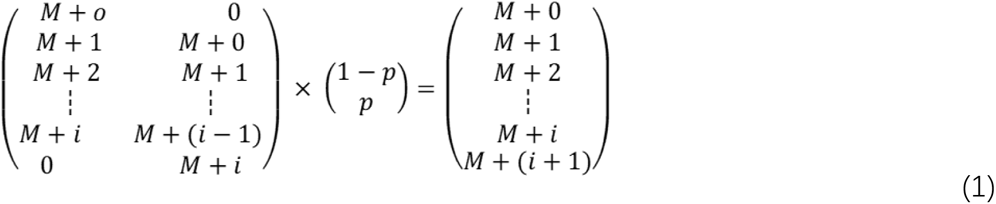

### Fatty acids and water-soluble metabolites labeling fraction calculation

Fatty acid features were extracted in EI-MAVEN software. The isotope natural abundance and impurity of labeled substrate were corrected using AccuCor written in R ^49^. Additionally, the labeling fraction was calculated automatically by the AccuCor.

### Statistics

Data were expressed as the mean ± SEM. Statistical analyses were conducted using GraphPad Prism version 9.1.0 or Microsoft Excel. Two-way ANOVA with Bonferroni post-test was performed to determine statistical significance for the time course study of tumor number and tumor burden. The log-rank test assessed the significance of the Kaplan-Meier analyses for progression-free survival. For comparisons between *G6pd^WT^;KL* and *G6pd^KO^;KL* lung tumors or cells in terms of metabolites, saponified fatty acids, and cell proliferation, a paired two-tailed Student’s t-test was employed. IHC quantification for Ki67, pS6, p-ERK, cleaved caspase3, p53, p21, 8-oxo-dG, and γ-H2AX was performed by obtaining representative images of entire lung lobes at each time point for each genotype or treatment, followed by statistical analysis using a paired two-tailed Student’s t-test. *P*<0.05 was considered statistically significant.

### Illustration tool

The schematic images are created using BioRender.com.

## Author Contributions

J.Y.G. was the lead principal investigator who conceived and supervised the project. J.Y.G. and T.L. designed the experiments, performed the data analysis, and interpreted the data. J.M.G. and J.D.R. provided *G6pd^Flox/Flox^* mouse strain, T.L., S.A., S.W., J.L., E.C.L., M.S., W.W., and V.B. performed most experiments. S.W. and M.S. performed IHC quantification. S.A., S.W., X.L., and J.L. maintained mouse colonies and mouse genotyping. Z.H. performed mouse surgery. C.L., X.S., J.D.R, and E.W. were involved in the project discussion. J.Y.G. and T.L. wrote the manuscript that was reviewed and edited by all authors.

## Acknowledgments

We are grateful to Eric Chiles and Yujue Wang in the Su laboratory for their assistance with LC-MS performance, and Jianming Wang in the Wenwei Hu laboratory for providing p53 and p21 antibodies for IHC. This work was supported by National Institute of Health (NIH) grant R01CA237347, American Cancer Society grant 134036-RSG-19-165-01-TBG, and Ludwig Princeton Branch of the Ludwig Institute for Cancer Research to J Guo; NIH grant R01CA163591 and Ludwig Princeton Branch of the Ludwig Institute for Cancer Research to E White; NIH grant P30 CA072720 to Rutgers Cancer Institute of New Jersey (Metabolomics Shared Resource, Biomedical Informatics shared resource, and Biospecimen Repository Service Shared Resource, at Rutgers Cancer Institute of New Jersey).

## Supplementary Methods

### *De novo* fatty acid synthesis analysis *in vitro*

*G6pd^WT^;KL* and *G6pd^KO^;KL* TDCLs were cultured in 6-cm dishes in RPMI medium 1640 without glucose (Gibco, #11879-020) supplemented with 10% fetal FBS, 1% Penicillin-Streptomycin, 0.075% sodium bicarbonate, and 2 g/L [U-^13^C_6_]-glucose for 24 hours and assessed in triplicate. Afterward, saponified fatty acids were extracted and subjected to LC-MS analysis for further analysis and calculation of ^13^C labeling fraction for fatty acids.

### Cultured cell sample preparation of saponified fatty acids for LC-MS analysis

Following the methodology described in a previous study ^41^, *G6pd^WT^;KL* and *G6pd^KO^;KL* TDCLs cultured in triplicate in 6-cm dishes were washed twice with PBS, followed by the addition of 1 mL of -20°C 90% methanol containing 0.3 mmol/L KOH. The resulting liquid, along with the cell debris, was scraped into 4 mL glass tubes. The samples were then heated at 80°C for 1 hour to saponify the fatty acids. After saponification, the samples were acidified with 100 µL of formic acid, followed by one minute of vortexing. The samples were extracted twice with 1 mL of hexane, and the organic phase was collected. The extracts were dried under air and dissolved in a mixture of methanol, chloroform, and isopropanol in a 1:1:1 ratio. All samples were vortexed for 20 seconds and then centrifuged at 4°C, 13,000 × g for 10 minutes. The resulting supernatant was transferred to LC-MS vials for subsequent analysis.

## Supplemental Figure Legend

**Figure S1.**
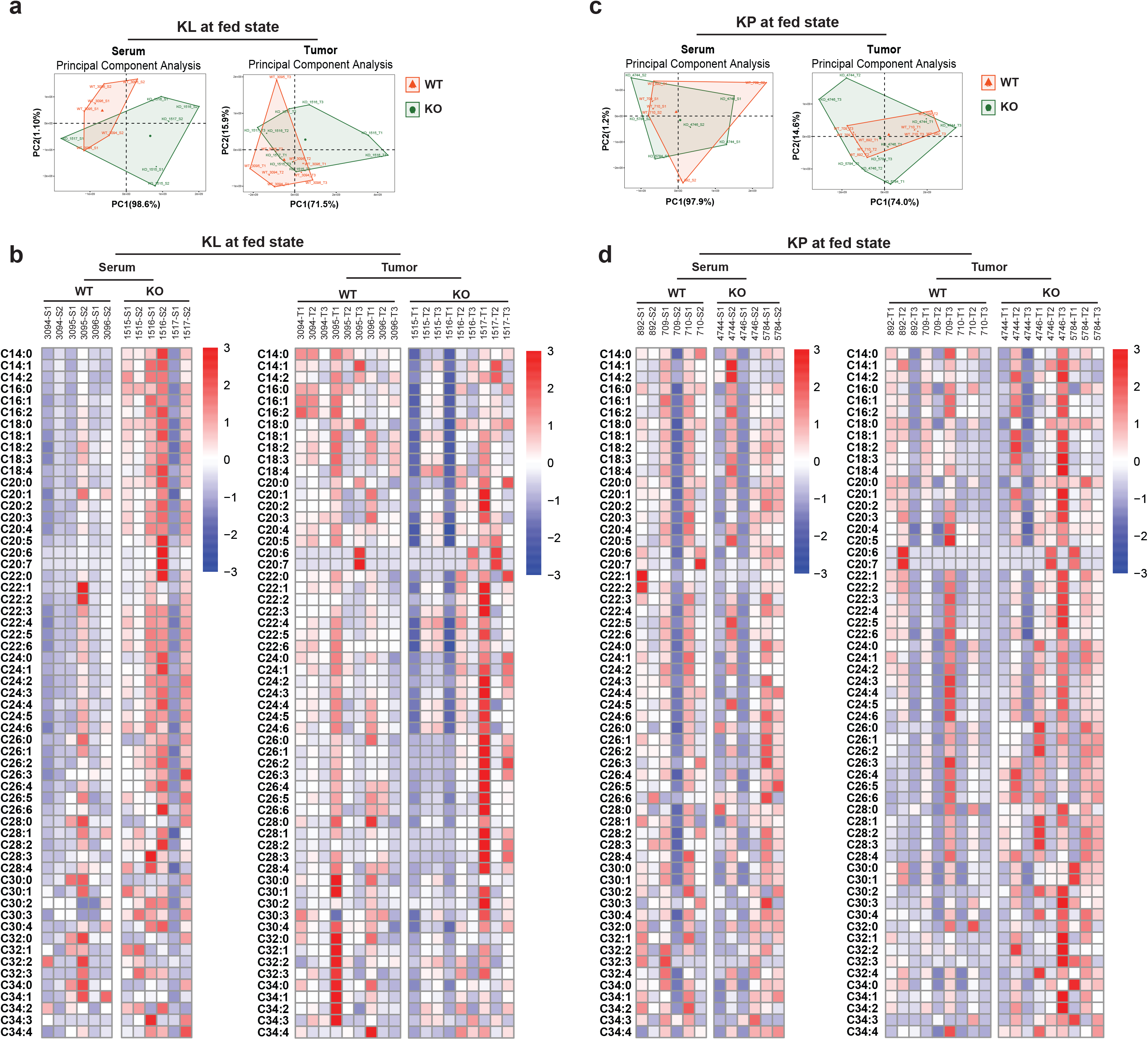
The pool size of fatty acids of serum and lung tumors. a & b. Principal Component Analysis (PCA) (a) and Heatmap (b) of saponified fatty acid pool size of *G6pd^WT^;KL* and *G6pd^KO^;KL* lung tumors and serum from KL tumor bearing mice in fed state at 12 weeks post-tumor induction. c & d. PCA (c) and Heatmap (d) of saponified fatty acid pool size of *G6pd^WT^;KP* and *G6pd^KO^;KP* lung tumors and serum from KP lung tumor bearing mice in fasted state (food was removed from the mice at approximately 9:00 a.m., and samples were collected at 3:00 p.m.) at 12 weeks post-tumor induction.

**Figure S2.**
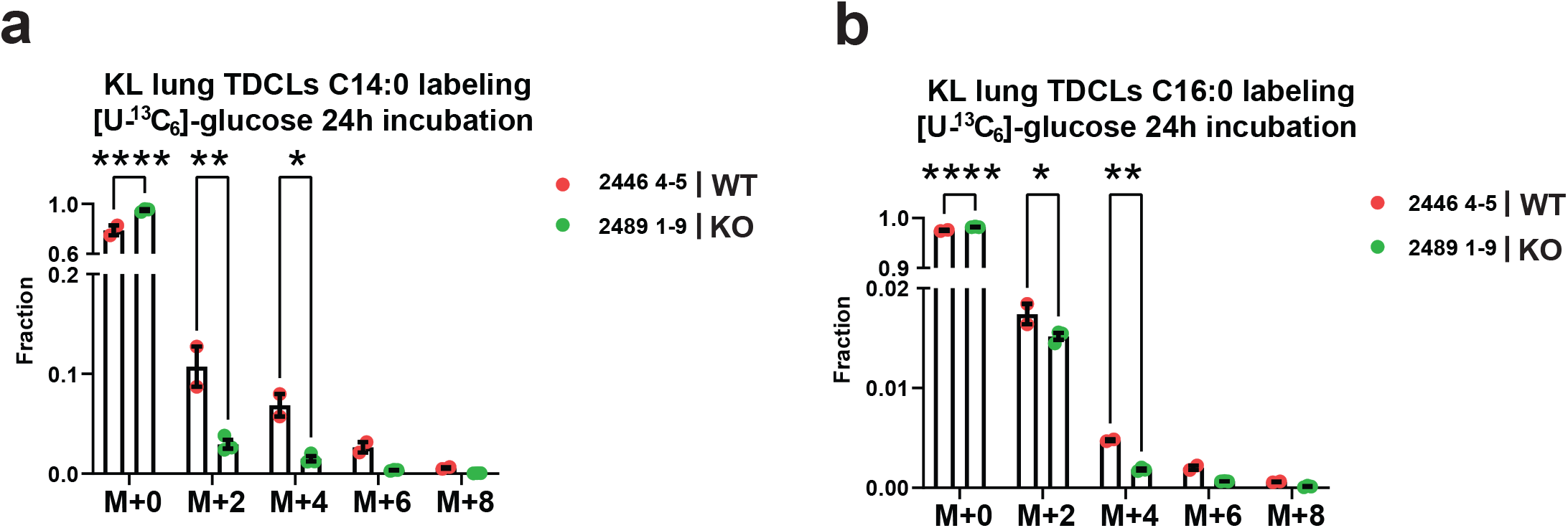
*De novo* fatty acid synthesis in KL lung TDCLs. a & b. C14:0 (a) and C16:0 (b) ^2^H labeling fraction of *G6pd^WT^;KL* and *G6pd^KO^;KL* TDCLs after 24 hours [U-^13^C_6_]-glucose labeling.

## Reference

1 Harrison, I. P. & Selemidis, S. Understanding the biology of reactive oxygen species and their link to cancer: NADPH oxidases as novel pharmacological targets. Clin Exp Pharmacol Physiol 41, 533–542, doi:10.1111/1440-1681.12238 (2014).

2 Ju, H. Q., Lin, J. F., Tian, T., Xie, D. & Xu, R. H. NADPH homeostasis in cancer: functions, mechanisms and therapeutic implications. Signal Transduct Target Ther 5, 231, doi:10.1038/s41392-020-00326-0 (2020).

3 Maddocks, O. D., Labuschagne, C. F. & Vousden, K. H. Localization of NADPH production: a wheel within a wheel. Molecular cell 55, 158–160, doi:10.1016/j.molcel.2014.07.001 (2014).

4 Goodman, R. P., Calvo, S. E. & Mootha, V. K. Spatiotemporal compartmentalization of hepatic NADH and NADPH metabolism. J Biol Chem 293, 7508–7516, doi:10.1074/jbc.TM117.000258 (2018).

5 Lewis, C. A. et al. Tracing compartmentalized NADPH metabolism in the cytosol and mitochondria of mammalian cells. Molecular cell 55, 253–263, doi:10.1016/j.molcel.2014.05.008 (2014).

6 Vander Heiden, M. G., Cantley, L. C. & Thompson, C. B. Understanding the Warburg effect: the metabolic requirements of cell proliferation. Science 324, 1029–1033, doi:10.1126/science.1160809 (2009).

7 Ding, H. et al. Activation of the NRF2 antioxidant program sensitizes tumors to G6PD inhibition. Sci Adv 7, eabk1023, doi:10.1126/sciadv.abk1023 (2021).

8 TeSlaa, T., Ralser, M., Fan, J. & Rabinowitz, J. D. The pentose phosphate pathway in health and disease. Nat Metab 5, 1275–1289, doi:10.1038/s42255-023-00863-2 (2023).

9 Cocco, P. Does G6PD deficiency protect against cancer? A critical review. J Epidemiol Community Health 41, 89–93, doi:10.1136/jech.41.2.89 (1987).

10 Pes, G. M., Bassotti, G. & Dore, M. P. Colorectal Cancer Mortality in Relation to Glucose - 6 - Phosphate Dehydrogenase Deficiency and Consanguinity in Sardinia: A Spatial Correlation Analysis. Asian Pac J Cancer Prev 18, 2403–2407, doi:10.22034/APJCP.2017.18.9.2403 (2017).

11 Pes, G. M., Errigo, A., Soro, S., Longo, N. P. & Dore, M. P. Glucose-6-phosphate dehydrogenase deficiency reduces susceptibility to cancer of endodermal origin. Acta Oncol 58, 1205–1211, doi:10.1080/0284186X.2019.1616815 (2019).

12 Dore, M. P., Davoli, A., Longo, N., Marras, G. & Pes, G. M. Glucose-6-phosphate dehydrogenase deficiency and risk of colorectal cancer in Northern Sardinia: A retrospective observational study. Medicine (Baltimore) 95, e5254, doi:10.1097/MD.0000000000005254 (2016).

13 Jiang, P. et al. p53 regulates biosynthesis through direct inactivation of glucose-6-phosphate dehydrogenase. Nat Cell Biol 13, 310–316, doi:10.1038/ncb2172 (2011).

14 Aurora, A. B. et al. Loss of glucose 6-phosphate dehydrogenase function increases oxidative stress and glutaminolysis in metastasizing melanoma cells. Proceedings of the National Academy of Sciences of the United States of America 119, doi:10.1073/pnas.2120617119 (2022).

15 Ghergurovich, J. M. et al. Glucose-6-phosphate dehydrogenase is not essential for K-Ras-driven tumor growth or metastasis. Cancer research, doi:10.1158/0008-5472.CAN-19-2486 (2020).

16 Shackelford, D. B. Unravelling the connection between metabolism and tumorigenesis through studies of the liver kinase B1 tumour suppressor. Journal of carcinogenesis 12, 16, doi:10.4103/1477-3163.116323 (2013).

17 Marcus, A. I. & Zhou, W. LKB1 regulated pathways in lung cancer invasion and metastasis. Journal of thoracic oncology : official publication of the International Association for the Study of Lung Cancer 5, 1883–1886, doi:10.1097/JTO.0b013e3181fbc28a (2010).

18 Shackelford, D. B. & Shaw, R. J. The LKB1-AMPK pathway: metabolism and growth control in tumour suppression. Nature reviews. Cancer 9, 563–575, doi:10.1038/nrc2676 (2009).

19 Garcia, D. & Shaw, R. J. AMPK: Mechanisms of Cellular Energy Sensing and Restoration of Metabolic Balance. Molecular cell 66, 789–800, doi:10.1016/j.molcel.2017.05.032 (2017).

20 Parker, S. J. et al. LKB1 promotes metabolic flexibility in response to energy stress. Metab Eng 43, 208–217, doi:10.1016/j.ymben.2016.12.010 (2017).

21 Jeon, S. M., Chandel, N. S. & Hay, N. AMPK regulates NADPH homeostasis to promote tumour cell survival during energy stress. Nature 485, 661–665, doi:10.1038/nature11066 (2012).

22 Skoulidis, F. et al. Co-occurring genomic alterations define major subsets of KRAS-mutant lung adenocarcinoma with distinct biology, immune profiles, and therapeutic vulnerabilities. Cancer discovery 5, 860–877, doi:10.1158/2159-8290.CD-14-1236 (2015).

23 Arbour, K. C. et al. Effects of Co-occurring Genomic Alterations on Outcomes in Patients with KRAS-Mutant Non-Small Cell Lung Cancer. Clinical cancer research : an official journal of the American Association for Cancer Research 24, 334–340, doi:10.1158/1078-0432.CCR-17-1841 (2018).

24 Li, S. et al. Assessing Therapeutic Efficacy of MEK Inhibition in a KRAS(G12C)-Driven Mouse Model of Lung Cancer. Clinical cancer research : an official journal of the American Association for Cancer Research 24, 4854–4864, doi:10.1158/1078-0432.CCR-17-3438 (2018).

25 Hasegawa, T. et al. Association Between the Efficacy of Pembrolizumab and Low STK11/LKB1 Expression in High-PD-L1-expressing Non-small-cell Lung Cancer. In Vivo 34, 2997–3003, doi:10.21873/invivo.12131 (2020).

26 Ngo, B., Van Riper, J. M., Cantley, L. C. & Yun, J. Targeting cancer vulnerabilities with high-dose vitamin C. Nature reviews. Cancer 19, 271–282, doi:10.1038/s41568-019-0135-7 (2019).

27 Hayes, J. D., Dinkova-Kostova, A. T. & Tew, K. D. Oxidative Stress in Cancer. Cancer cell 38, 167–197, doi:10.1016/j.ccell.2020.06.001 (2020).

28 Lee, W. N. et al. Measurement of fractional lipid synthesis using deuterated water (2H2O) and mass isotopomer analysis. Am J Physiol 266, E372–383, doi:10.1152/ajpendo.1994.266.3.E372 (1994).

29 Hellerstein, M. K. et al. Measurement of de novo hepatic lipogenesis in humans using stable isotopes. The Journal of clinical investigation 87, 1841–1852, doi:10.1172/JCI115206 (1991).

30 Zhang, Z. et al. Serine catabolism generates liver NADPH and supports hepatic lipogenesis. Nat Metab 3, 1608–1620, doi:10.1038/s42255-021-00487-4 (2021).

31 Zhang, Z., Chen, L., Liu, L., Su, X. & Rabinowitz, J. D. Chemical Basis for Deuterium Labeling of Fat and NADPH. J Am Chem Soc 139, 14368–14371, doi:10.1021/jacs.7b08012 (2017).

32 Niu, X. et al. Cytosolic and mitochondrial NADPH fluxes are independently regulated. Nat Chem Biol, doi:10.1038/s41589-023-01283-9 (2023).

33 Ciccarese, F., Zulato, E. & Indraccolo, S. LKB1/AMPK Pathway and Drug Response in Cancer: A Therapeutic Perspective. Oxid Med Cell Longev 2019, 8730816, doi:10.1155/2019/8730816 (2019).

34 Svensson, R. U. et al. Inhibition of acetyl-CoA carboxylase suppresses fatty acid synthesis and tumor growth of non-small-cell lung cancer in preclinical models. Nat Med 22, 1108–1119, doi:10.1038/nm.4181 (2016).

35 Cararo-Lopes, E. et al. Autophagy buffers Ras-induced genotoxic stress enabling malignant transformation in keratinocytes primed by human papillomavirus. Cell Death Dis 12, 194, doi:10.1038/s41419-021-03476-3 (2021).

36 Eriksson, S. E., Ceder, S., Bykov, V. J. N. & Wiman, K. G. p53 as a hub in cellular redox regulation and therapeutic target in cancer. Journal of molecular cell biology 11, 330–341, doi:10.1093/jmcb/mjz005 (2019).

37 Chen, L. et al. NADPH production by the oxidative pentose-phosphate pathway supports folate metabolism. Nat Metab 1, 404–415 (2019).

38 Song, J., Sun, H., Zhang, S. & Shan, C. The Multiple Roles of Glucose-6-Phosphate Dehydrogenase in Tumorigenesis and Cancer Chemoresistance. Life (Basel) 12, doi:10.3390/life12020271 (2022).

39 Mele, L. et al. A new inhibitor of glucose-6-phosphate dehydrogenase blocks pentose phosphate pathway and suppresses malignant proliferation and metastasis in vivo. Cell Death Dis 9, 572, doi:10.1038/s41419-018-0635-5 (2018).

40 Cerami, E. et al. The cBio cancer genomics portal: an open platform for exploring multidimensional cancer genomics data. Cancer discovery 2, 401–404, doi:10.1158/2159-8290.CD-12-0095 (2012).

41 Bhatt, V. et al. Autophagy modulates lipid metabolism to maintain metabolic flexibility for Lkb1-deficient Kras-driven lung tumorigenesis. Genes & development 33, 150–165, doi:10.1101/gad.320481.118 (2019).

42 Guo, J. Y. et al. Autophagy suppresses progression of K-ras-induced lung tumors to oncocytomas and maintains lipid homeostasis. Genes & development 27, 1447–1461, doi:10.1101/gad.219642.113 (2013).

43 Khayati, K. et al. Autophagy compensates for Lkb1 loss to maintain adult mice homeostasis and survival. Elife 9, doi:10.7554/eLife.62377 (2020).

44 Khayati, K. et al. Transient Systemic Autophagy Inhibition is Selectively and Irreversibly Deleterious to Lung Cancer. Cancer research, doi:10.1158/0008-5472.CAN-22-1039 (2022).

45 Zhang, Z. et al. Serine catabolism generates liver NADPH and supports hepatic lipogenesis. Nat Metab, doi:10.1038/s42255-021-00487-4 (2021).

46 Hui, S. et al. Glucose feeds the TCA cycle via circulating lactate. Nature, doi:10.1038/nature24057 (2017).

47 Tsugawa, H. et al. MS-DIAL: data-independent MS/MS deconvolution for comprehensive metabolome analysis. Nat Methods 12, 523–526, doi:10.1038/nmeth.3393 (2015).

48 Tsugawa, H. et al. A lipidome atlas in MS-DIAL 4. Nat Biotechnol 38, 1159–1163, doi:10.1038/s41587-020-0531-2 (2020).

49 Su, X., Lu, W. & Rabinowitz, J. D. Metabolite Spectral Accuracy on Orbitraps. Anal Chem 89, 5940–5948, doi:10.1021/acs.analchem.7b00396 (2017).

